# Mecp2 deletion results in profound alterations of developmental and adult functional connectivity

**DOI:** 10.1101/2021.10.28.466323

**Authors:** Rachel M. Rahn, Allen Yen, Siyu Chen, Seana H. Gaines, Annie R. Bice, Lindsey M. Brier, Raylynn G. Swift, LeiLani Lee, Susan E. Maloney, Joseph P. Culver, Joseph D. Dougherty

## Abstract

As a regressive neurodevelopmental disorder with a well-established genetic cause, Rett Syndrome and its *Mecp2* loss-of-function mouse model provide an excellent opportunity to define potentially translatable functional signatures of disease progression, as well as offer insight into *Mecp2*’s role in functional circuit development. Thus, we applied optical fluorescence imaging to assess mesoscale calcium functional connectivity (FC) in the Mecp2 cortex prior to symptom onset as well as during decline. We found that FC was profoundly disrupted in Mecp2 males both in juvenile development and early adulthood. Female Mecp2 mice displayed a subtle homotopic contralateral increase in motor cortex as juveniles but not in adulthood, where instead parietal regions were implicated. Additionally, conditional rescue studies indicated FC phenotypes are driven by excitatory neurons. Altogether, the female results identify subtle candidate translatable biomarkers of disease progression, while the male results indicate MeCP2 protein is needed in a circuit-specific manner for FC.

## Introduction

Rett Syndrome (RTT) is a monogenic regressive neurodevelopmental disorder (NDD) with clearly-defined symptomatology but differences in individual presentation and progression that present a challenge in management and treatment of the disorder. RTT is an X-linked disorder caused by mutations in the methyl-CpG-binding protein 2 (*MeCP2*) gene and is known for its strong sex-specific effects, with RTT occuring in approximately 1 in every 10,000 to 15,000 live female births while male RTT patients are relatively rare^1^. Progression of the disease occurs over a series of stages, with development appearing to proceed normally for the first 6-18 months of life, followed by the first disease stage which may include decreasing head growth and abnormal muscle tone^2, 3^. A period of regression, including the development of stereotypical hand wringing movements and a loss of language and purposeful hand use, is then followed by a stabilization where some behavior may improve^1, 3^. However, symptoms such as ataxia and motor issues often develop or persist, along with seizures and respiratory abnormalities^1–3^. The last stage includes a motor deterioration with reduced mobility^2^. With improved interventions to manage seizure and aspiration risks, mean lifespan in RTT patients has increased, yet survival rates are still much reduced compared to the general population^4–6^. In light of the increased mortality and severity of physical symptoms that RTT patients develop, a variety of clinical trials of therapeutics have been launched that aim to ameliorate or reverse the condition. However, limited understanding of the underlying biological mechanisms creates challenges in knowing how timing and targeting of interventions may affect outcomes. Therefore, characterizing RTT-related *Mecp2* phenotypes in clinically relevant modalities such as functional neuroimaging could help shed light on the circuits perturbed during disease progression.

Furthermore, although the molecular consequences of MECP2 loss are well-characterized, the subsequent circuit pathobiology is less clear. Alterations to the excitatory-inhibitory balance have been documented in the Mecp2 mouse and is hypothesized to be a driving force behind many RTT-related deficits^7, 8^. Aberrations during critical periods also are speculated to cause long-term effects to the many circuits affected by RTT. The precocious onset and closure of the visual critical period in *Mecp2*-null mice appears to be related to a previously reported early maturation of GABAergic neurons^9^. As RTT is an X-linked disorder, there is a strong difference between the Mecp2 sexes in mice: Mecp2 females are less affected than males due to mosaic expression of the mutant and wildtype alleles caused by X-inactivation, just as seen in the human population. In mice, Mecp2 males typically die between 7 and 17 weeks of age, with symptom onset as early as 4 weeks, while females do not die rapidly from the disorder and have a much delayed symptom onset at 16 weeks or later^10^. As males model a faster, more severe decline ending in death, they have often been used to characterize phenotypes that relate to the clinical symptoms at earlier ages and with greater effect. Mecp2 females, meanwhile, more directly relate to the primarily-female clinical population and so are often used to detect more subtle versions of the male phenotypes, in the hope of translating findings and treatments to female RTT patients. While the *Mecp2* mouse model has been heavily studied for its gene expression and cellular deficits, mesoscale connectivity and the flow of information between brain regions is understudied. Previous work on visual and auditory-evoked potentials demonstrate that abnormal responses occur in RTT^11, 12^, and studies that focus on quantitative definition of RTT phenotypes have identified potential EEG biomarkers of RTT disease state^13–15^. However, compared to physiological and synaptic or molecular phenotypes, RTT functional neuroimaging and other mesoscale phenotypes still remain relatively unexplored. Mecp2 mice therefore are a powerful tool by which to identify patterns of normalized brain function that are relevant to RTT patients and the understanding of the effects of developmental *Mecp2* dysfunction. The role of *Mecp2* in development as an epigenetic regulator of synaptic maturation is well-established^16^, but additional roles of the gene are continuing to be explored, suggesting that additional circuits may play a role in RTT progression. Therefore it is essential to further explore how *Mecp2* dysfunction affects global and localized function in the *in vivo* mouse brain during development and adulthood.

Functional connectivity (FC) neuroimaging provides a non-invasive method by which phenotypes and potential biomarkers for NDDs such as RTT can be established. RTT imaging has previously been largely confined to structural measures, including reported decreases in both gray and white matter volume^17, 18^. However, structural changes are not expected to be especially dynamic and thus would likely reflect less change in the disease state across time. In contrast, functional imaging does respond acutely to changes in the brain’s ongoing tasks, and chronically in response to a variety of manipulations and therapeutics^19, 20^. Of the functional imaging paradigms, resting-state FC is relatively efficient, mapping all of the connections between all regions-of-interest within the field-of-view simultaneously. Further, resting state FC does not require task compliance, an advantage when evaluating NDD populations. FC also has potential as an early marker of disease state that will become evident later in development, as previous work has suggested that FC at 6 months of age is predictive of autism spectrum disorder (ASD) diagnosis at 2 years amongst infants at a high risk for ASD^21^. However, establishing RTT imaging signatures in a clinical population is challenging due to the heterogeneity of genetics and environment intrinsic to such a group. Identifying early FC phenotypes is additionally complicated by the fact that diagnosis often is in secondary stages of the disorders and therefore the earliest time points prior to significant decline are missed.

Mouse models of NDDs such as RTT could serve as a powerful tool by which to establish FC-based disease phenotypes. Inbred mouse lines such as the *Mecp2* loss-of-function mouse model of RTT permit an increased control of genetics and environmental factors, so that the effects of a single disease-related manipulation can be isolated. The faster maturation of the mouse also facilitates the imaging of disease-related decline without waiting for the years required in the clinical population. In addition, mouse models allow for a potentially larger sample size to detect more subtle phenotypes, which is particularly essential in disorders such as RTT with a low incidence rate in the general population. Finally, the Mecp2 mouse also permits mechanistic studies to be performed and circuit- and molecular-level manipulations. Rescue of *Mecp2* expression throughout the adult brain using the Mecp2^lox-Stop^ system results in dramatic rescue of RTT-like phenotypes in mice^10^, suggesting that therapies introduced late in life are viable for correcting the disorder. Furthermore, rescue of MeCP2 specifically in GABAergic neurons has been demonstrated to significantly improve male lifespan as well as characteristic behavioral deficits^22^. We have focused here on the GABAergic rescue rather than the brainwide rescue, as we presumed it would provide the most specific information regarding which, of any, normalized functional connections correspond to an amelioration of the disease. Previous studies have documented RTT-related alterations to the excitation-inhibition balance; research focused on inhibitory signaling in RTT indicates that this imbalance is at least partially be driven by interneuron-related dysfunction^22, 23^. However, other studies have indicated that excitation alterations may also occur in parallel to these inhibitory changes^7, 24^. The question of the exact cell populations that contribute to the excitation-inhibition imbalance observed in RTT therefore continues to be an area of interest in the field, and our rescue in GABAergic populations will investigate whether inhibition is the primary driver behind any Mecp2 FC phenotype found. The ability to use genetic tools like the Mecp2^lox-Stop^ mouse to rescue MeCP2 levels in certain cellular subpopulations therefore provides a powerful opportunity to identify how FC parallels partial rescue of protein levels in RTT, and test the hypothesis that early FC changes may predate behavioral alterations that have previously been demonstrated to emerge during adulthood in Mecp2 females.

To better understand if FC might be a viable biomarker for *Mecp2*-related disease progression, we used widefield optical fluorescence imaging to characterize cortical FC in both Mecp2 males and females at both a developmental time point as well as a time point adjacent to the canonical symptom onset and decline period in each sex (Figure 1A). We discovered FC abnormalities in affected Mecp2 mice of both sexes, with a more severe phenotype appearing in males as expected. However, to our surprise, an interneuron-specific protein-level rescue did not exhibit a behavioral rescue, in contrast to prior work^22^, suggesting subtle differences between the two vGAT Cre lines might present an opportunity to finemap survival-relevant cells. FC traits were also mostly unrescued, although males displayed an intermediate homotopic connectivity phenotype. Nonetheless, we were able to establish that motor FC abnormalities appear early in both sexes and exhibit promise as detectable phenotypes that mirror underlying circuit perturbations and the complex biological role of *Mecp2* before significant decline occurs.

**Figure 1:**
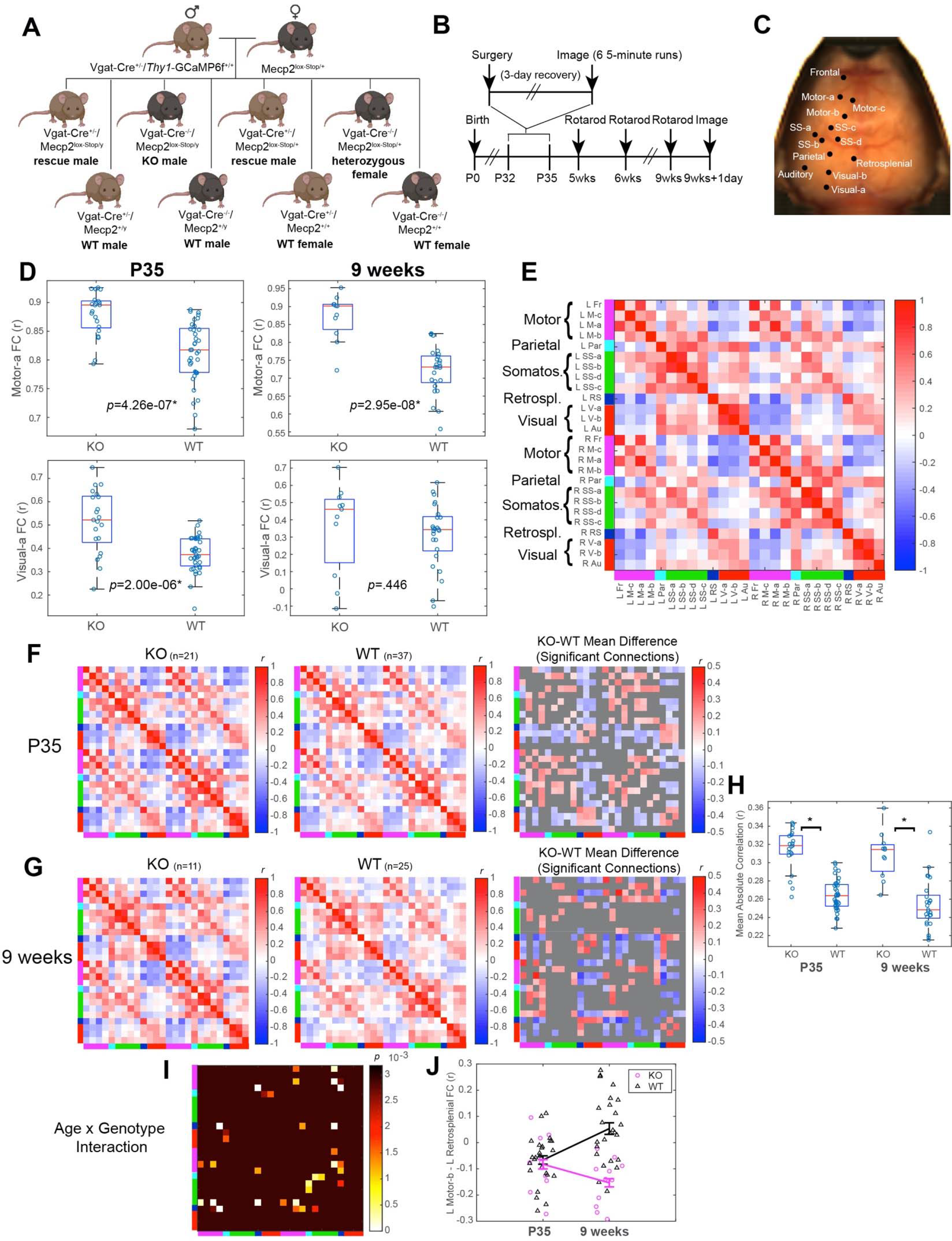
Mecp2 KO males display multiple differences compared to controls at both P35 and 9 weeks of age. A) Breeding a Mecp2 female and male Vgat-Cre/*Thy1*-GCaMP6 mouse produced Mecp2 affected, GABAergic neuron-specific rescue, and WT animal cohorts. (Graphic created using BioRender.com.) B) Timeline of imaging and behavioral data collection from Mecp2 male mice. C) Representative brain image displaying the positions of the 13 cortical seeds in the left hemisphere. (Remaining 13 of 26 total cortical seeds are positioned identically in the right hemisphere.) D) Homotopic contralateral comparisons between the Motor-a seed pair at P35 and 9 weeks (row 1) and between the Visual-a seed pair at P35 and 9 weeks (row 2). E) Representative matrix figure displaying the seed names and composition of the seed-by-seed FC matrices reflecting correlation coefficients (Pearson r) for each seed pair. Seed abbreviations: L left, R right, Fr Frontal, M-c Motor 3rd seed, M-a Motor 1st seed, M-b 3rd seed, Par Parietal, SS-a Somatosensory 1st seed, SS-b Somatosensory 2nd seed, SS-d Somatosensory 4th seed, SS-c Somatosensory 3rd seed, RS Retrosplenial, V-a Visual 1st seed, V-b Visual 2nd seed, Au Auditory. F) P35 and G) 9-week group means of Mecp2 KO males (1st column), Mecp2 WT controls (2nd column), and significant differences (Pearson r) between the two groups after an FDR correction of an unpaired t-test (3rd column). H) Differences in the average absolute strength of FC across each mouse’s cortex (P35 *p=*3.37e-13*, 9 weeks *p=*2.32e-06*). Red lines are group medians, box bounds represent the 25th (bottom) and 75th (top) percentiles of the group, and each circle represents one mouse. I) Significant age-by-genotype interactions after FDR comparison of rmANOVA. J) Scatter plot of the connection strength between left Motor-b and left Retrosplenial seeds for each mouse with data at both P35 and 9 weeks of age. Lines connect the group means at each time point with error bars representing the standard error of the mean (SEM) (0.4-4.0 Hz).

## Results

### Mecp2 males display widespread FC abnormalities at both P35 and 9 weeks of age

Mecp2 males were imaged during development at postnatal day (P)35 and at 9 weeks, during the model’s typical decline period, with rotarod motor coordination collected adjacent to each time point (Figure 1B). Male Mecp2 mice traditionally display a more severe phenotype than females, reflecting the effects of complete MeCP2 loss in males compared to mosaic loss of MeCP2 expression in heterozygous females. As symptom onset in Mecp2 males can be as early as 4 weeks^25^, we hypothesized that we would observe a significant FC deficit at P35. Specifically, we predicted that we would observe an overconnectivity phenotype in the visual cortex that reflects previous functional studies focused specifically in this region^9^. We also predicted we would observe a significant difference from controls in motor regions due to the documented Mecp2 gait and motor coordination deficits^22, 26^. We therefore first tested for our hypothesized differences in bilateral Motor-a and Visual-a seed connections by comparing the correlations of calcium signals across time between homotopic seed pairs in these regions. We observed a significant increase in FC in the bilateral Motor- a connections at both time points, and between left and right Visual-a seeds at P35 but not at 9 weeks (Figure 1C-D).

Following these initial analyses, we performed a discovery-based analysis of all 26 cortical seeds’ connections with each other. Visual cortex’s FC with other regions such as somatosensory and motor cortex were significantly different between the Mecp2 knock-out (KO) and wildtype (WT) males at both time points (Figure 1E-G, Figure S1). Motor-motor FC was significantly different in multiple seed pairs at 9 weeks but not P35, paralleling the worsening of motor deficits previously reported in the literature. We noticed that across all connections, positively correlated regions had stronger correlations in the KOs as compared to WT controls, while the anti-correlated regions were more negatively correlated in the KOs at both time points. To assess this impression empirically, we subsequently tested whether the absolute value of each functional connection, averaged together across the cortex to produce one value per mouse, was significantly different between groups. At both ages, KO mice had significantly higher mean absolute FC than WT controls (Figure 1H).

To also directly evaluate potential functional correlates of decline, we examined differences between groups with age, by performing an age-by-genotype rmANOVA comparison. Multiple connections were found to have a significant age-by-genotype interaction (Figure 1H), particularly in retrosplenial cortical regions (Figure 1I). The connection between left Motor-b and left Retrosplenial seeds displayed the most significant interaction in males (*p*=1.74e-05), exhibiting a reduction in KO connection strength while WT males’ FC increased between P35 and 9 weeks of age (Figure 1J).

Beyond seed-seed FC analyses, we also performed a pixel-by-pixel homotopical contralateral FC analysis, in order to investigate whether there were aberrations in callosal connectivity. By examining each pixel’s connectivity with its homotopic equivalent in the other cortex, we examined bilateral relationships not dependent on our 26 canonical functional seeds’ placement. We found that Mecp2 KO males displayed bilateral overconnectivity in motor and visual cortices at P35 and motor overconnectivity at 9 weeks, although the posterior region overconnectivity at this age appeared to be more concentrated in retrosplenial than visual cortex (Figure 2). Significant age-by-genotype interactions were observed in anterior cortex (Figure 2D), while bilateral connectivity as represented by a single mean FC value for each mouse displayed a decrease in both KO and WT males across time (Figure 2E).

**Figure 2:**
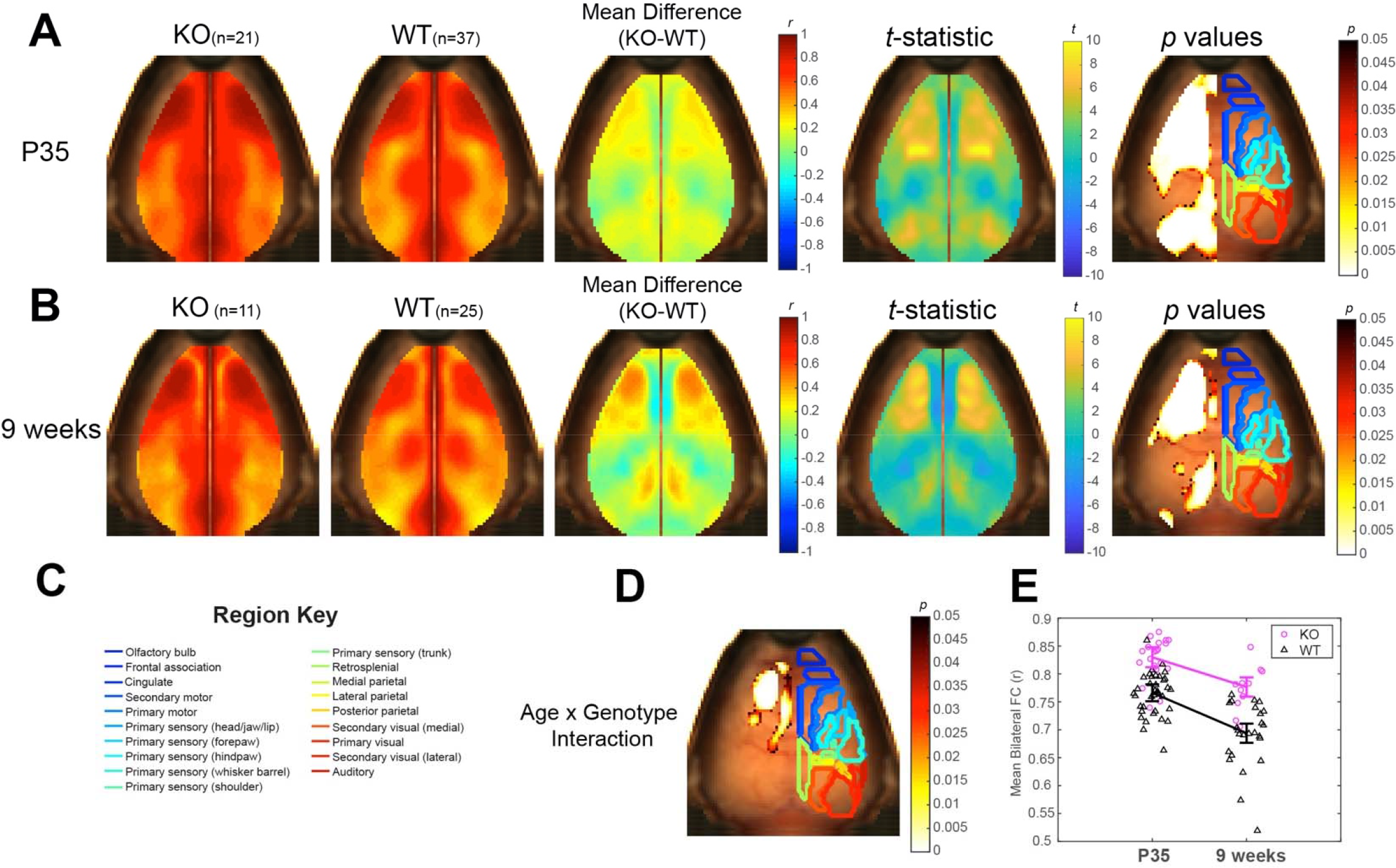
Mecp2 KO males display bilateral FC alterations compared to controls at both P35 and 9 weeks of age. A) P35 and B) 9-week group means of Mecp2 KO (1st column) and WT (2nd column) bilateral pixel maps, with difference maps (3rd column), *t*- statistic maps (4th column) and regions of significant change after a cluster-based correction (5th column), with significant *p*-values on the left hemisphere and Paxinos atlas region boundaries outlined on the right, overlaid on a background brain image. C) Paxinos atlas-based region outlines mapped on a representative dorsal cortical surface. D) Significant age-by-genotype interaction derived from rmANOVA. E) Scatter plot of the mean bilateral connection strength of all pixels significant in D for each mouse sampled at both P35 and 9 weeks of age. Lines connect the group means at each time point with error bars representing SEM (0.4-4.0 Hz).

While cortical seed measures and specifically our placement have previously been demonstrated to have high between-mouse accuracy^27^, seed-based measures do depend upon accuracy of functional region boundaries. To further examine connectivity in the cortex independent of cortical seed placement, we therefore performed node degree analyses^19^ to determine if the number of strong positive connections that each pixel (node) exhibits with other areas of the brain changes in Mecp2 males. At both P35 and 9 weeks of age, Mecp2 males displayed a strong increase in node degree in anterior regions that include frontal and motor cortices, as well as the majority of visual and retrosplenial cortices (Figure 3). A significant age-by-genotype interaction was observed in visual and retrosplenial regions (Figure 3D); overall, the average node degree of each pixel across the cortex declined in both Mecp2 males and WTs across development (Figure 3E). The increase in Mecp2 males’ node degree mirrored the similar increase in bilateral connectivity at both ages that we previously reported (Figure 2), reflecting an overall increase in the strength of the strongly correlated connections, a group of connections that typically includes bilateral measures because of homotopic contralateral regions’ functional similarity.

**Figure 3:**
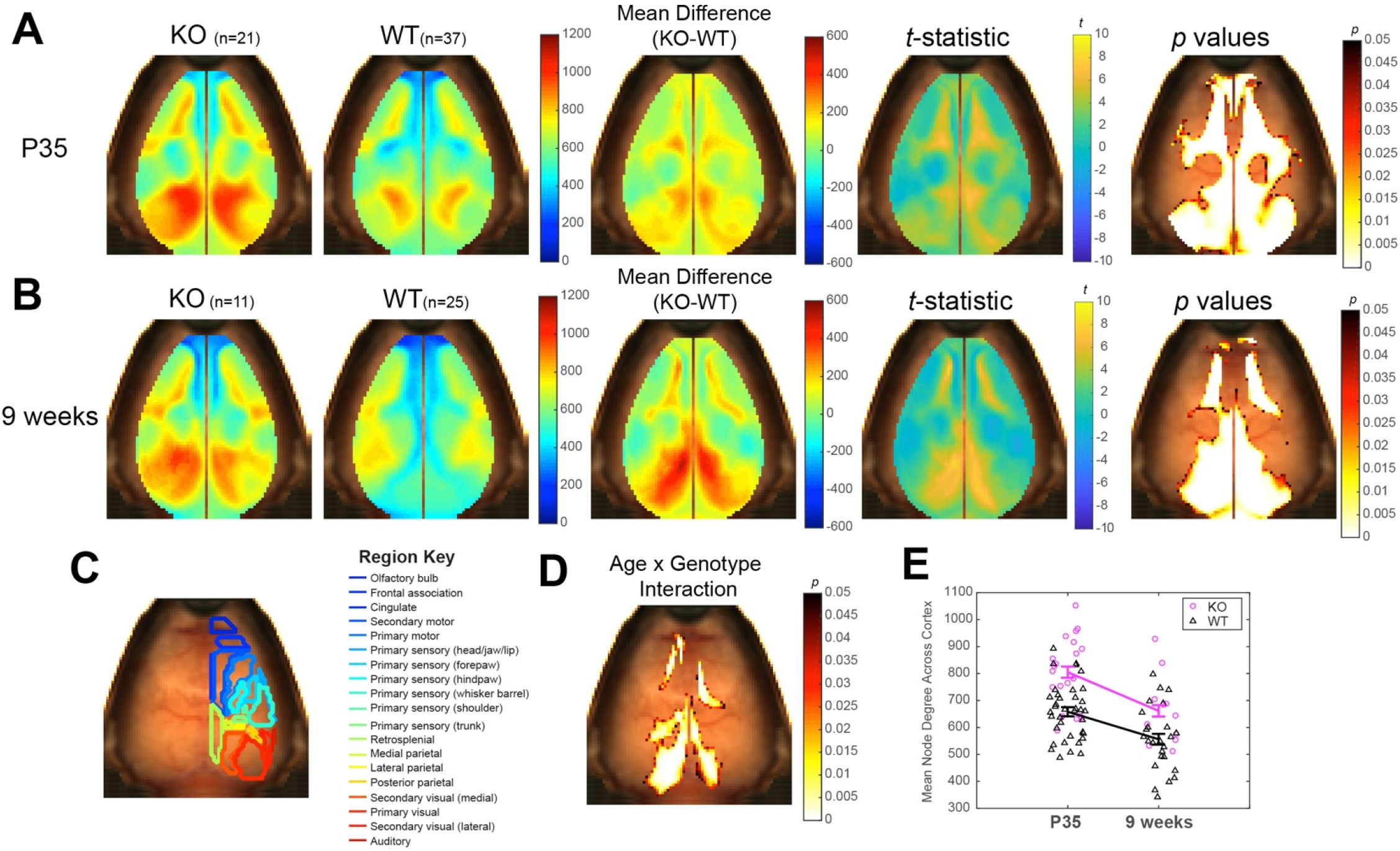
Mecp2 KO males display increased node degree compared to controls at both P35 and 9 weeks of age. A) P35 and B) 9-week group means of Mecp2 KO (1st column) and WT (2nd column) node degree maps, with difference maps (3rd column), *t*- statistic maps (4th column) and regions of significant change after a cluster-based correction (5th column). C) Paxinos atlas-based region outlines mapped on representative dorsal cortical surface. D) Significant age-by-genotype interaction derived from rmANOVA. E) Scatter plot of the mean node degree for the average of all significant pixels in D for each mouse with data at both P35 and 9 weeks of age. Lines connect the group means at each time point with error bars representing SEM (0.4-4.0 Hz).

### Mecp2 females display alterations in homotopic contralateral FC and node degree

Because RTT is an X-linked disorder, the Mecp2 model displays less severe behavior deficits and symptoms in females than males. The early imaging time point we used at P35 is prior to symptom onset reported in the literature^10^, and we therefore hypothesized that any observed FC phenotype would be subtle, if present. We also hypothesized that motor FC deficits would thus appear later at 4 months of age and a similar visual FC phenotype would appear as we hypothesized in males, because of previous reports of a precocious visual critical period^9^. However, imaging results indicated that there was no significant change in seed-seed FC in either these targeted functional regions at either P35 or 4 weeks nor after performing a wider discovery-based analysis (Figure 4A-E). The changes in absolute connectivity observed in complete MeCP2 loss (males, Figure 1H), were not observed in females (Figure 4E), indicating that expression of MeCP2 in at least half of all neurons was sufficient to rescue normal mesoscale connection strength. However, a significant age-by-genotype interaction occurred between left Auditory and right Motor-a seed-seed as well as right Parietal and right Visual-b connection (Figure 4F). Examining the more significant connection (L Aud-R Motor-a) in detail revealed an increase in FC in WT females with aging which did not occur in het females (Figure 4G). Additionally, an increase in homotopic contralateral FC occurred at P35 near the border between motor and somatosensory cortices (Figure 5), subtly mirroring our observed overconnectivity phenotypes in males. At 4 months of age, posterior regions overlapping with the parietal cortex and anterior regions including motor cortex displayed significantly reduced bilateral connectivity in females (Figure 5B). No pixels survived a multiple comparison correction to have a significant age-by-genotype interaction (Figure 5D), but mean bilateral connectivity across the entire cortex did display a mean increase in Mecp2 het females compared to WTs at P35, then a mean decrease compared to WTs at 4 months (Figure 5E). Altogether, females displayed a limited age-by-genotype interaction in analysis of a 325-connection set, while bilateral measures displayed differences between genotypes that differed in direction dependent on the age sampled.

**Figure 4:**
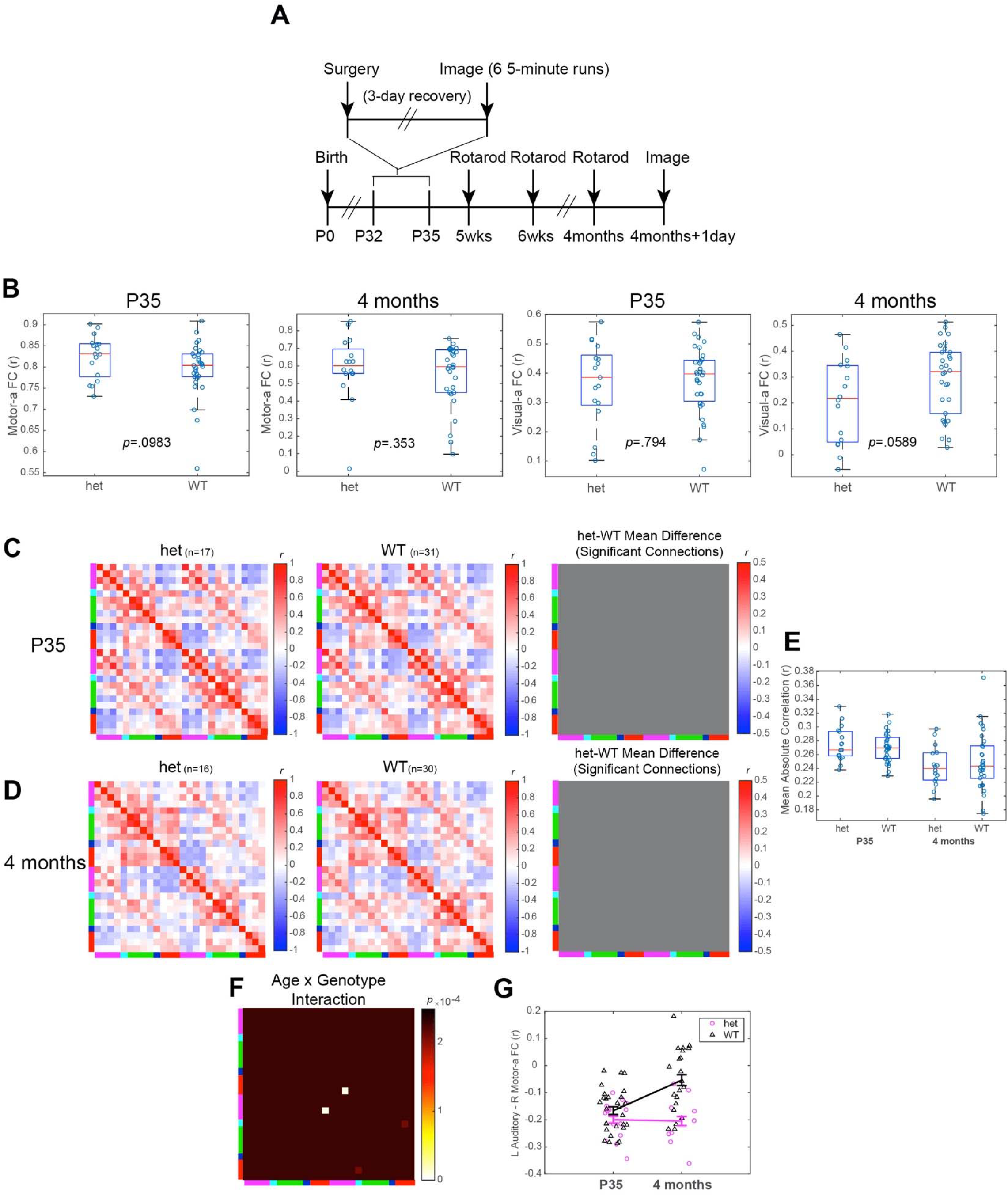
Mecp2 heterozygous females and WT controls display a time-by-genotype interaction but no significant differences from controls at each individual time point. A) Timeline of imaging and behavioral data collection from Mecp2 female mice. B) Homotopic contralateral comparisons between the Motor-a seed pair at P35 (column 1) and 4 months (column 2) and between the Visual-a seed pair at P35 (column 3) and 4 months (column 4). C) P35 and D) 4-month group means of Mecp2 het females (1st column), Mecp2 WT controls (2nd column), and significant differences (Pearson r) between the two groups after an FDR correction of an unpaired t-test (3rd column). E) Differences in the average absolute strength of FC across each mouse’s cortex (P35 *p=*0.488, 4 months *p=*0.530) (0.4-4.0 Hz). Red lines are group medians, box bounds represent the 25th (bottom) and 75th (top) percentiles of the group, and each circle represents one mouse. F) Significant age-by-genotype interactions after FDR comparison of rmANOVA. G) Scatter plot of the connection strength between left Motor-b and left Retrosplenial seeds for each mouse with data at both P35 and 4 months of age. Lines connect the group means at each time point with error bars representing SEM (0.4-4.0 Hz).

**Figure 5:**
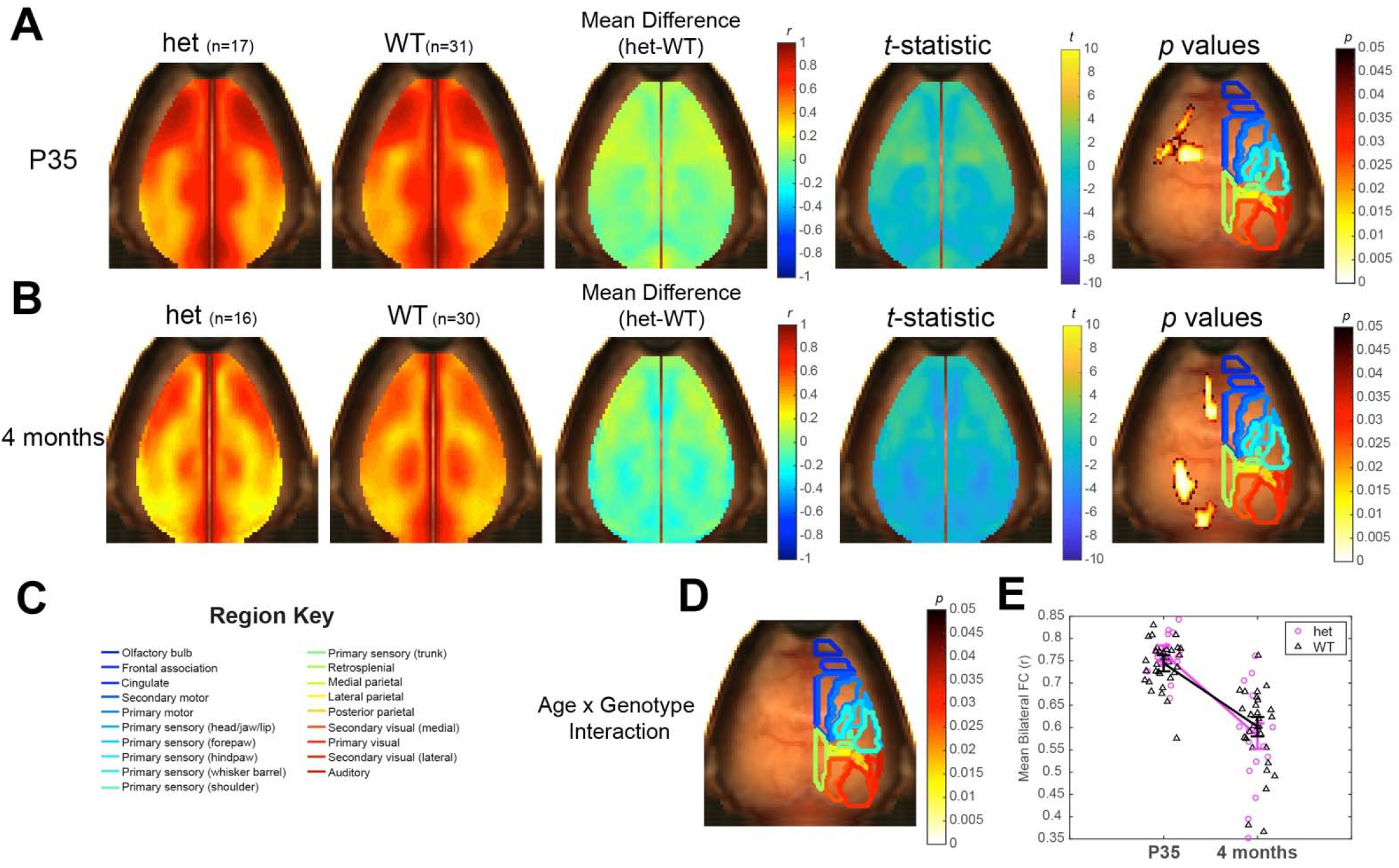
Mecp2 heterozygous females display age-dependent bilateral FC alterations compared to controls. A) P35 and B) 4- month group means of Mecp2 het (1st column) and WT (2nd column) bilateral pixel maps, with difference maps (3rd column), *t*- statistic maps (4th column) and regions of significant change after a cluster-based correction (5th column), with significant *p*-values on the left hemisphere and Paxinos atlas region boundaries outlined on the right, overlaid on a background brain image. C) Paxinos atlas-based region outlines mapped on representative dorsal cortical surface. D) Significant age-by-genotype interaction derived from rmANOVA. E) Scatter plot of the mean bilateral connection strength of all cortical pixels. These pixels are averaged to create a single value for each mouse with data at both P35 and 4 months of age. Lines connect the group means at each time point with error bars representing SEM (0.4-4.0 Hz).

Beyond seed-based measures of regional correlation, we next tested whether Mecp2 disease state might be reflected in other measures of connectivity. We found that node degree, a measure of strong positive connectivity, was also perturbed in Mecp2 females. Node degree differences between Mecp2 females and controls reflected the traditional worsening of symptoms between the time points sampled, as no significant differences were observed at P35 but a decrease in node degree was detected at 4 months (Figure 6A-B). This direction of change and its location mirror the bilateral underconnectivity also observed in Mecp2 females at 4 months (Figure 5B) but has only limited spatial overlap with the decreased node degree that Mecp2 males exhibited compared to controls (Figure 3). A significant age-by-genotype interaction occurred in the right posterior cortex and in a limited section of the anterior cortex (Figure 6C-D). Mean node degree across the cortex also appeared overall to have a steeper decline between ages in Mecp2 females than WTs (Figure 6E). Altogether, node degree was affected later in development, at 4 months of age, than bilateral measures, but additionally displayed an age-by-genotype interaction as Mecp2 females exhibited a steeper decline in the measure.

**Figure 6:**
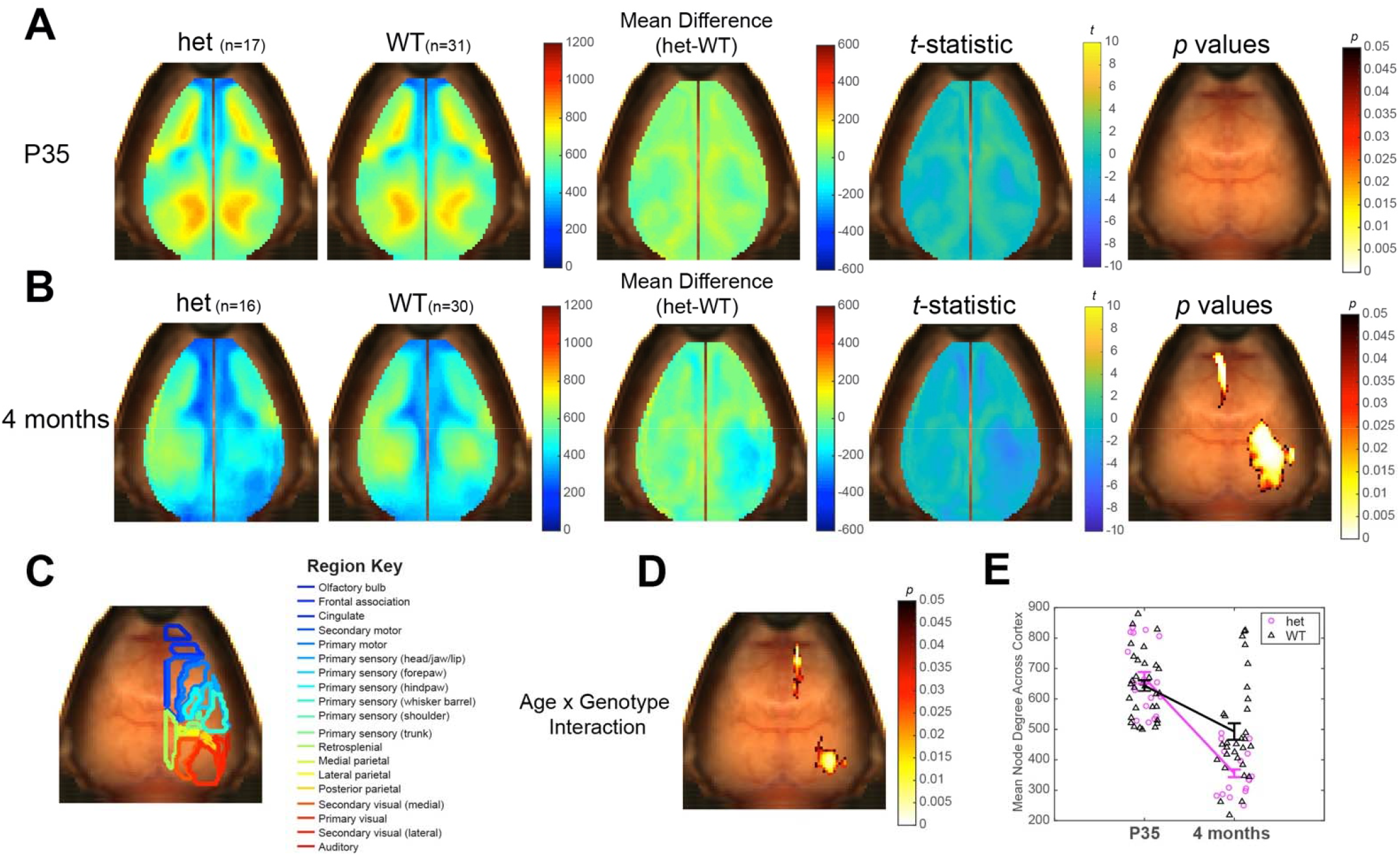
Mecp2 heterozygous females display decreased node degree compared to controls at 4 months of age. A) P35 and B) 4-month group means of Mecp2 het (1st column) and WT (2nd column) node degree maps, with difference maps (3rd column), *t*- statistic maps (4th column) and regions of significant change after a cluster-based correction (5th column). C) Paxinos atlas-based regions outlined mapped on representative dorsal cortical surface. D) Significant age-by-genotype interaction derived from rmANOVA. E) Scatter plot of the mean node degree for the average pixel of all significant pixels in D for each mouse with data at both P35 and 4 months of age. Lines connect the group means at each time point with error bars representing SEM (0.4-4.0 Hz).

### Interneuron-specific MeCP2 rescue fails to ameliorate FC deficits in females and only partially rescues male phenotype

Based on previous research that documented a rescue in MeCP2 in a GABAergic interneuron-specific population that improves survival and rotarod motor performance^22^, we also tested if a GABAergic interneuron-specific rescue would rescue the FC phenotype. We used a well-validated mouse line expressing Cre protein under the control of an IRES sequence knocked into the endogenous locus of the vesicular GABA transporter (vGAT). This was the same promoter utilized in previous work, though used in a BAC context by the prior studies^22^. vGAT Cre mice were crossed to a genetically inducible MeCP2 rescue line in which Cre mediates replacement of a key mutant exon with a healthy wildtype version. Though the Cre line is well validated as specific and comprehensive for GABAergic neurons, each floxed allele can have different efficiencies^28^. We therefore first confirmed that the Cre comprehensively rescued *Mecp2* expression in GABAergic neurons of the mouse brain. We used immunofluorescence to confirm the expected 50% reduction of expression of *Mecp2* in heterozygous female mice and the complete knock-out in KO males (Figure S3). Further, MeCP2 rescue in GABAergic neurons displayed robust restoration of expression throughout the brain (Figure S4). To quantify the extent of this rescue, we counted the number of MeCP2+ GABAergic neurons, NeuN+ neurons, and non-neuronal DAPI+-only cells in the striatum, cortex, and cerebellum (Figures S3-7). We focused on the cortex because our imaging technique queries that region, and the striatum because a large portion of cells would be expected to be GABAergic, and both striatum and cerebellum are involved in motor function. These experiments confirm *Mecp2* expression is restored in this key subpopulation of neurons throughout the brain, in expected proportions.

However, to our surprise, we saw only one significant difference between the rescued Mecp2 and unrescued mice, either male or female, in assessed behavioral and survival phenotypes (Figure 7A-B). While both males and females displayed a main effect of genotype on rotarod performance when rescued, unrescued, and WT mice were evaluated across time (male *p=*9.98e-05, females *p=*4.33e-06), post hoc testing showed only a significant difference between rescued and unrescued animals at 4 months of age in females (*p=*0.0196). For all other phenotypes, both survival metrics and rotarod performance, the genetically rescued animals were not distinguishable from unrescued animals, in contrast to prior work. In addition, we observed a largely parallel lack of rescue in FC phenotype (Figure 7C-D). Crucially, we did observe one change upon rescue: a subtly intermediate bilateral FC phenotype in males, greater than controls but less than unrescued Mecp2 FC strength (Figure 7E-H). However, the rescue animals were otherwise indistinguishable from unrescued mice in all FC analyses (Figures 7 & 8). Further, we observed that both had similar significant differences when compared to controls. Thus, while we had hoped to identify brain circuits that corresponded to behavioral rescue, the lack of behavioral rescue precluded this analysis. However, we could confirm that FC also did not ameliorate, demonstrating some consistency between FC and behavioral traits.

**Figure 7:**
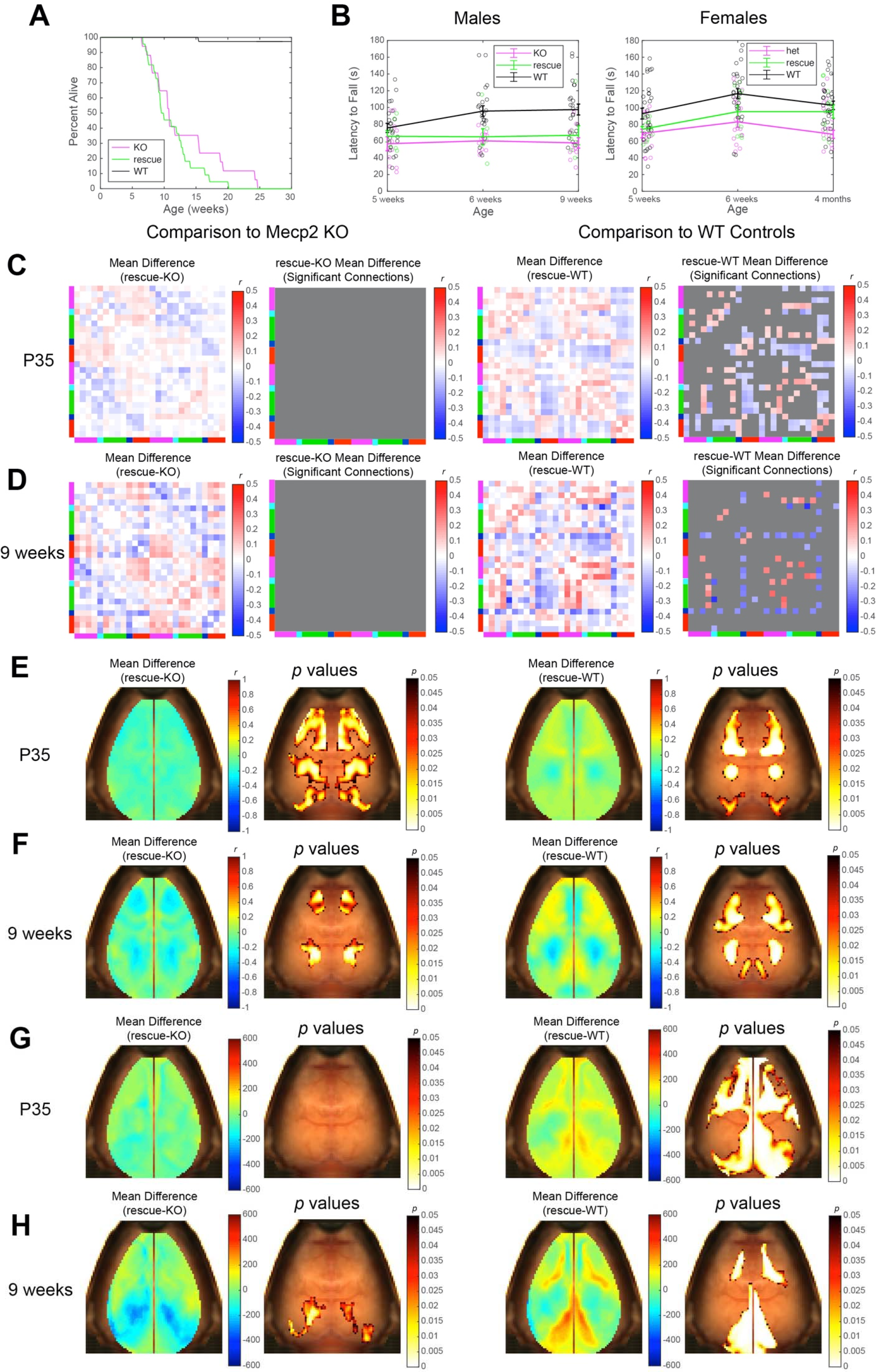
Behavior and survival were not rescued as expected after GABAergic *Mecp2* rescue, with a very limited FC rescue observed in males. A) Survival curve for Mecp2 male mice across time (Mecp2 males n=17, rescues n=22, WT n=37). B) Mean accelerating Rotarod latency to fall in males (left) and females (right) across time and scatterplot of each individual’s latency averaged between two runs (Mecp2 males n=11, rescues n=7, WT n=23; Mecp2 females n=16, rescues n=12, WT n=29). C) Left: P35 difference between Mecp2 rescue and unrescued KO males (column 1 difference matrix, column 2 FDR-corrected significant connections after t-test comparison). Right: P35 difference between Mecp2 rescue and WT controls (column 3 difference matrix, column 4 FDR-corrected significant connections after t-test comparison). D) Left: 9-week difference between Mecp2 rescue and unrescued KO males (column 1 difference matrix, column 2 FDR-corrected significant connections after t-test comparison). Right: Nine-week difference between Mecp2 rescue and WT controls (column 3 difference matrix, column 4 FDR-corrected significant connections after t-test comparison). E) P35 bilateral pixel difference map and regions of significant change between (left) Mecp2 rescue and unrescued KO males, and between (right) Mecp2 rescue and WT control males. F) 9-week bilateral pixel difference map and regions of significant change between (left) Mecp2 rescue and unrescued KO males, and between (right) Mecp2 rescue and WT control males. G) Left: P35 node degree difference map and regions of significant change between (left) Mecp2 rescue and unrescued KO males, and between (right) Mecp2 rescue and WT control males. H) 9-week node degree difference map and regions of significant change between (left) Mecp2 rescue and unrescued KO males, and between (right) Mecp2 rescue and WT control males. All pixel-wise significance maps represent results after performance of a cluster-based correction (P35 KO n=21, rescue n=22, WT n=37; 9 weeks KO n=11, rescue n=7, WT n=25) (0.4-4.0 Hz).

**Figure 8:**
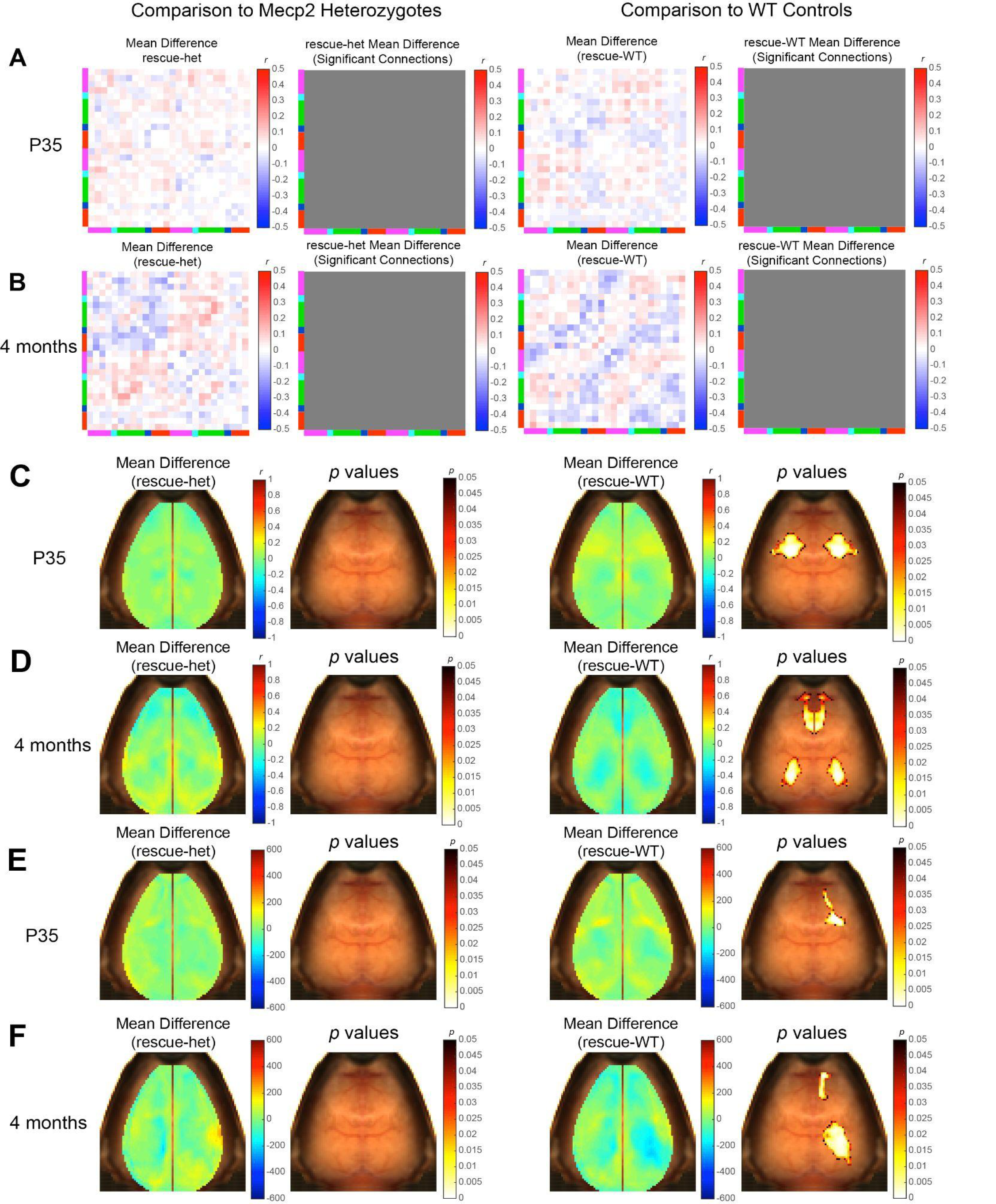
GABAergic-specific rescue females did not display a rescue in FC compared to unrescued Mecp2 females. A) Left: P35 difference between the GABAergic-specific Mecp2 rescue and unrescued Mecp2 het females (column 1 difference matrix, column 2 FDR-corrected significant connections after t-test comparison. Right: P35 difference between Mecp2 rescue and WT controls (column 3 difference matrix, column 4 FDR-corrected significant connections after t-test comparison. B) Left: 4-month difference between Mecp2 rescue and unrescued Mecp2 het females (column 1 difference matrix, column 2 FDR-corrected significant connections after t-test comparison). Right: 4-month difference between Mecp2 rescue and WT controls (column 3 difference matrix, column 4 FDR-corrected significant connections after t-test comparison). C) P35 bilateral pixel difference map and regions of significant change between (left) Mecp2 rescue and unrescued Mecp2 het females, and between (right) Mecp2 rescue and WT controls. D) 4-month bilateral pixel difference map and regions of significant change between (left) Mecp2 rescue and unrescued Mecp2 het females, and between (right) Mecp2 rescue and WT controls. E) P35 node degree difference map and regions of significant change between (left) Mecp2 rescue and unrescued Mecp2 het females, and between (right) Mecp2 rescue and WT controls. F) 4-month node degree difference map and regions of significant change between (left) Mecp2 rescue and unrescued Mecp2 het females, and between (right) Mecp2 rescue and WT controls. All pixel-wise significance maps represent results after performance of a cluster-based correction (P35 heterozygous n=17, 15 rescue n=15, WT n=31; 4 months heterozygous n=16, rescue n=14, WT n=30) (0.4-4.0 Hz).

**Table 1.** Mecp2 deletion causes extensive effects in males and limited effects in females in both development and adulthood. Size of arrows indicates size of effect, with a large arrow reflecting more than 50% of connections or pixels affected or a single *t*-test *p* value less than 0.001, a medium arrow reflecting more than 25% affected or a single *p* value less than 0.01, and small arrow less than 25% affected or single significant *p* value greater than 0.01.

| IMAGING<br>PARAMETER | MALES P35 | MALES<br>ADULT | FEMALES P35 | FEMALES<br>ADULT |
| --- | --- | --- | --- | --- |
| Mutant Effects Compared to WT Controls |  |  |  |  |
| Bilateral Motor<br>FC | ↑ | ↑ | - |  |
| Bilateral Visual<br>FC | ↑ | - | - |  |
| Overall Seed-Seed FC | ↑↓ | ↑↓ | - | - |
| Absolute FC<br>Amplitude | ↑ | ↑ | - | - |
| Bilateral<br>Pixelwise FC | ↑ | ↑ | ↑ | ↓ |
| Node Degree | ↑ | ↑ | - | ↓ |
| Rescue Effects Compared to Mutants |  |  |  |  |
| Overall Seed-Seed FC | - | - | - | - |
| Bilateral<br>Pixelwise FC | ↓ | ↓ | - | - |
| Node Degree | - | ↓ | - | - |
| Rescue Effects Compared to WT Controls |  |  |  |  |
| Overall Seed-Seed FC | ↑↓ | ↑↓ | - | - |
| Bilateral<br>Pixelwise FC | ↑↓ | ↑↓ | ↑ | ↓ |
| Node Degree | ↑ | ↑ | ↑ | ↓ |

## Discussion

By examining sex-specific effects in the Mecp2 mouse model of Rett Syndrome both during development and proximal to symptom onset, we have established that Mecp2 mice exhibit both developmental and adult FC abnormalities (Table 1). These FC abnormalities are more evident in males: Mecp2 males display a variety of FC abnormalities at both P35 and 9 weeks of age, with extensive overconnectivity in motor and visual FC at both time point, including visual alterations at P35, which is proximal to the visual critical period. The female deficits observed seem to be largely limited to later underconnectivity in multiple measures in posterior regions such as the parietal cortex. In regard to early aberrations, females display homotopic contralateral overconnectivity in the motor cortex similar to what was already seen in males and much earlier than the canonical onset of symptoms in the Mecp2 model, establishing that a developmental FC increase is evident in both sexes.

As seed-seed FC abnormalities in Mecp2 males displayed both increases and decreases depending upon functional region, these changes reflected an overall increase in absolute connection strength. This included an overconnectivity phenotype in males’ bilateral FC and node degree at both ages, and in females’ bilateral FC at P35. This increase in FC strength is consistent with prior overconnectivity hypotheses for other NDDs such as autism spectrum disorder (ASD), where overconnectivity in a variety of networks has been observed in clinical populations^29^. Although the direction of change in ASD appears to be highly dependent on sex, brain region, age, and sampling methods, increased FC has been previously reported, for example, in the sensorimotor network in boys with ASD^30^, and it is possible that RTT and ASD may share commonalities in an FC phenotype in certain networks. The overconnectivity we observed in Mecp2 mice might also be a result of a shift in the excitation/inhibition balance, which has been previously observed to change in Mecp2 mice^31^. Reduced inhibition in the connections we analyzed might be the cause of the overconnectivity observed, although some prior studies would suggest the opposite effect due to a shift in the balance to favor inhibition over excitation^32, 33^. Mecp2 mice have previously been shown to have a precocious visual critical period^9^, which might explain why a P35 overconnectivity phenotype is observed in the visual cortex. Although alterations in motor FC were predicted, an underconnectivity phenotype was anticipated as motor task performance deteriorated. The motor overconnectivity that we detected, although not in the expected direction of effect, might affect motor behaviors in other ways and impede information processing between motor regions. Bilateral motor connectivity increases may additionally negatively affect connections between motor and association cortices necessary to perform complex motor tasks. There are also significant declines in FC between certain motor seeds and select visual and somatosensory regions at both time points, suggesting that associations between non-homotopic brain regions may break down as motor performance declines in the Mecp2 model.

In regard to sex-specific effects, the difference between the prominence and presence of a FC phenotype in males and females reflects human RTT symptomatology and indicates that future experiments may need to image females at even older ages. The Mecp2 mouse model previously has been documented as having symptom onset anywhere between 4 and 14 months in females^10, 25^, so more subtle FC differences may not appear until mice age further (though 3 months post-windowing surgery approaches the limit for our cranial windows’ lifespan, suggesting a cross sectional approach might be needed). While the P35 female bilateral phenotype echoes the overconnectivity seen in males at both ages, the significantly perturbed area shifts to also include posterior regions by 4 months in the females, where underconnectivity is observed similar to the decrease in node degree also observed in 4-month females. Node degree alterations in Mecp2 females at this older age are also in the opposite direction of effect as male node degree perturbations, as KO males consistently display higher connectivity than controls. This change in direction of effect importantly suggests that just as seen in our analysis of the males’ set of 325 seed-seed comparisons, Mecp2 aberrations do not solely reflect an increase in connectivity or a global state change, such as the altered arousal state previously observed in Mecp2 mice^34^. This suggests that just as RTT clinical populations transition through disease stages, so too might the Mecp2 female mouse, which reflects the sex of the majority of RTT individuals, progress through discrete stages of disease progression. These changes begin with a motor overconnectivity that precedes the later coordination deficits characteristic of this model. Subsequently, the differences between Mecp2 and control females move on to become parietal-based FC deficits in adulthood after motor FC alterations resolve. Additional behavior tests probing individual differences may better stratify imaging data and detect correlated or predictive FC signatures that shed light on sympatology we did not measure here.

The lack of rescue of the FC phenotype when we performed a GABAergic interneuron-specific rescue (using Vgat, also known as Viaat, Cre) was unexpected but paralleled a general lack of behavioral and lifespan rescue. Although there was a significant post hoc difference in rotarod performance at 4 months in females, that was the only behavioral change observed between rescued and unrescued animals, and the female FC phenotype failed to rescue while the more severely affected males demonstrated a total lack of rescue in survival, behavioral or FC metrics. This lack of rescue in males, and in female FC, shed light on the cell populations key to a rescue. First and foremost, we learned that inhibitory signaling is not the predictor of FC deficits, and instead the excitation within the excitation-inhibition balance is the meaningful driver for our measure. Specifically, the gross functional abnormalities still present after we rescued MeCP2 in inhibitory neurons, and their only very limited difference from unrescued animals demonstrates that FC deficits primarily reflect differences in excitatory neurons. The fact that the females’ more subtle motor phenotype displayed some improvement on rotarod while the males did not also brings up the intriguing possibility that there may be a sex-specific effect in the role of neuronal subpopulations in the Mecp2 phenotype.

As to why male lifespan was not rescued, we suspect the difference between our findings and the prior studies of a rescue following a GABAergic rescue may reflect either Cre line or background strain differences. First, we considered the potential for strain differences, as we had performed more backcrosses to C57Bl/6J mice than the aforementioned study. Strain differences might also provide an opportunity, if the rescue was in fact mediated by an FVB allele in tight linkage with the BAC insertion site. However, the use of the same BAC mouse to also delete *Mecp2* and cause motor and survival deficits largely rules this out^23^. Our study also used the Vgat-ires-Cre which is a slightly different line than the bacterial artificial chromosome (BAC) Viaat-Cre line previously used in the literature, although both are supposed to target the same neuronal subpopulation, using the same gene promoter. The line we used, as a knock-in, should be the most complete recapitulation of endogenous Viaat expression, though it is possible the IRES dampens Cre expression to a level where recombination is not complete. We therefore next examined this possibility. After our initial characterization (Figures S3-4), given the lifespan findings we conducted a series of additional immunofluorescence studies (Figures S5-7), including colabeling with GAD67, and largely ruled out this explanation, as we saw the expected recombination in all brain regions examined (Figures S4-7), and total recombinant cell numbers are in line with the distribution and expected number of vGAT neurons from published in situ hybridization data^35, 36^ and other reported expression patterns^37^.

BAC line Cres and Cres made by knock-in to the endogenous genes can differ in Cre expression for other reasons as well. BACs can be at a higher copy number, potentially leading to a higher Cre expression level and thus different recombination rate, perhaps in cells that only transiently express Cre during development. In addition, though it is more rare with BACs than smaller transgenes, locus of integration effects, or loss of distal enhancer elements, can alter the pattern of expression allowing for some ectopic targeting. Therefore, if additional regions displayed recombination with the BAC beyond those seen in the knock-in, those would be high priority for mediating the rescue (rather than VGAT neurons broadly as was previously concluded). Indeed, there may be an opportunity to map the key cell type down substantially by careful comparison of these two lines to identify any cells distinct between the two, if the BAC mice were to be rederived from their frozen stocks. We would note that our data so far already provide the benefit of ruling out vast swaths of GABAergic neurons as being responsible for the survival and motor rescue: GABA cells from leading regions for motor behavior such as cortex, striatum and cerebellum can all be ruled out as not being sufficient to normalize behavior and thus our findings might guide thinking about gene therapy approaches. We suspect the key cells that differ between the BAC and the knock-in line may be found in the brainstem or spinal cord based on a study showing survival rescue in a posterior Cre (HoxB1)^38^, and early lethality due to MECP2 deletion below cervical level 2 (Cdx2 Cre)^39^. Indeed, if it is possible to rescue survival with MECP2 re-expression in just a small cell population in a discrete brain region, then gene therapy approaches might be more trackable.

Overall, our study demonstrated that extensive FC abnormalities occur in male Mecp2 mice and that subtle motor-related FC deficits are also evident in females at P35, well before what is typically considered the detectable symptom onset. Motor FC phenotypes might thus deserve examination as a possible measure of altered development in the clinical population as well and serving as an early sign of functional recovery in RTT gene therapy or other therapeutics.

## Acknowledgments

This work was supported by the National Institute of Neurological Disorders and Stroke (F31NS110222, R.M.R, R01NS099429, J.P.C. and R01NS090874, J.P.C.), National Institute of Mental Health (R01MH124808 and R01MH107515, J.D.D.), and National Institute of Child Health and Human Development (P50HD103525, Intellectual and Developmental Disabilities Research Center at Washington University).

## Methods

### Animal Care and Use

All studies were performed under the guidelines established by the Washington University Institutional Animal Care and Use Committee and according to the approved animal protocol. All mice were housed in a facility with 12:12 hour light-dark cycle with water and food freely available. A total of 143 mice (21 Vgat-Cre^-/-^/Mecp2^lox-Stop/y^ (KO) males, 22 Vgat-Cre^+/-^/Mecp2^lox-Stop/y^ (rescue) males, 37 Vgat-Cre^+/-^/Mecp2^+/y^ or Vgat-Cre^-/-^/Mecp2^+/y^ (WT) males, 17 Vgat-Cre^-/-^/Mecp2^lox-Stop/+^ (heterozygous) females, 15 Vgat-Cre^+/-^/Mecp2^lox-Stop/+^ (rescue) females, and 31 Vgat-Cre^-/-^/Mecp2^+/+^ or Vgat-Cre^-/-^/Mecp2^+/+^ (WT) females, all carrying a *Thy1*/GCaMP6f allele) were used for the P35 imaging and behavior analysis performed (Figure 1A). A subset of these mice, those who were alive at the decline time point and passed data quality thresholds, were used for the decline and longitudinal analyses at 9 weeks (males, KO n=11, rescue n=7, WT n=25) and 4 months (females) (heterozygous n=16, rescue n=14, WT n=30). To generate this cohort, female Mecp2^lox-Stop/+^ mice were bred to male mice homozygous for *Thy1*/GCaMP6f (JAX:024276) and heterozygous for the Vgat-Cre allele (JAX:016962)^45^. Because Mecp2 mutants are often poor mothers, following birth each litter was transferred within 0-2 days to a lactating female CD1-IGS (Charles River:022) who had also given birth within 2 days of the experimental litter, and Mecp2 litters were raised by the CD1-IGS foster female with two of her own CD1-IGS/C57Bl6J pups until weaning. This fostering scheme was based upon the Mecp2 fostering protocol described by Vogel Ciernia, et al^46^. All litters consisted of 4-8 pups, as all litters larger than 8 pups were culled to reduce potential maternal care confounds.

### Rotarod and Bird Scoring Behavior

Rotarod data were collected one day prior to imaging, at 5- and 6-week timepoints with 7 days between data collection days, and at 9 weeks for surviving males and 4 months for surviving females. On each testing day, mice were habituated to the testing room for 30 minutes and then run on the rotarod for 5 trials: trial 1 was a 60-second stationary trial, trials 2 and 3 were 60-second continuous trials run at 3 revolutions/sec, and trials 4 and 5 were 180-second accelerating trials that began at 3 revolutions/sec and accelerated at +1.0 revolutions/sec. The time each mouse fell off the rod was noted and averaged for the two continuous trials and two accelerating trials to derive a single continuous and single accelerating value. Following completion of the rotarod assay, a Bird score of Mecp2 symptom severity was performed, in which mobility, gait, hindlimb clasping, tremor, breathing, and general condition are marked on a 0-2 scale to track disease progression^10, 47^.

Due to facility access restrictions during to the COVID-19 pandemic, 5 mice (4 males, 1 female) lacked 6-week data, and 5 mice (3 males, 2 females) had a 7-week time point collected instead of 6-week; all were excluded from behavioral data analysis. Approximately 30 mice also had weekly rotarod data collected past the 6-week timepoint before pandemic restrictions necessitated the temporary cessation of behavior data and the restructuring of data collection to only coincide with imaging time points. Males lacking 9-week data either died before that time point due to complications from their RTT-like phenotype or could not be tested due to COVID-19 laboratory access restrictions; these mice were excluded from behavioral analysis.

### Optical Fluorescence Imaging

All mice had their scalp retracted under isoflurane anesthesia and a chronic optical window made of Plexiglas adhered to the dorsal cranial surface using dental cement (C&B-Metabond, Parkell Inc., Edgewood, NY) at postnatal day (P)32 in order to facilitate optical imaging during wakefulness. Three days post-surgery, mice were evaluated using the Bird score of Mecp2 symptom progression and then imaged in a widefield optical fluorescence imaging system ^48^ allowing for simultaneous calcium fluorescence and hemodynamic data collection. Four LEDs (470 nm (GCaMPf excitation), 530 nm, 590 nm, and 625 nm) shone on the cortex sequentially at a rate of 16.8 Hz per LED, and data were collected at the same rate using a sCMOS camera. Field of view was approximately 1.1 cm^2^ and 78x78 pixels after on-camera binning and a spatial downsample. The data were also temporally downsampled to 8.4 Hz before analysis was completed. Imaging was performed on P35 for all mice and again at 9 weeks of age for males and 4 months for females. All mice had 30 minutes of resting-state data collected during wakefulness in each imaging session, which comprised 6 runs of 5 minutes each.

### Immunofluorescence

Mice were perfused between 8 and 10 weeks of age using 4% paraformaldehyde and the brains were extracted. Following an overnight fixation, brains were cryoprotected with a 15% and 30% (w/v) sucrose solution, cryosectioned into 30-μm slices, and stored at 4°C in PBS/0.1% Azide until use. Free-floating sections were washed with PBS, permeabilized in 0.1% Triton X-100 and blocked with 5% normal donkey serum in PBS, then incubated with primary antibodies (1:1000 chicken anti-GFP (Aves GFP-1020), 1:200 rabbit anti-MeCP2 (D4F3) XP (Cell Signaling 3456S), and 1:500 mouse anti-GAD67 (Millipore MAB5406)) at 4°C overnight. The next day, sections were washed with PBS and then incubated for one hour at room temperature with secondary antibodies (1:1000 donkey anti-chicken AF488, 1:1000 donkey anti-rabbit AF568, and 1:1000 donkey anti-mouse AF647). Nuclei were stained blue with DAPI, and sections were mounted with ProLong Gold (ThermoFisher P36934).

Immunofluorescence images were captured using a Zeiss AxioScan Z1 or Zeiss LSM700 confocal microscope with a Plan-Neofluar 10x/0.30 objective. The 405nm, 488nm, 555nm, and 639nm laser lines were used to visualize DAPI, Alexa Fluor488, Alexa Fluor568, and Alexa Fluor647, respectively. Pseudocolor images and cell counts were processed using ImageJ.

### Data Quantification and Analysis

All analyses were performed in MATLAB (2017b). Data processing and baseline analyses were based on the processing pipeline documented in Brier et al^49^. All runs with a variance in raw light levels greater than 1.5% were discarded and not used in analysis. Mice were additionally screened to confirm they displayed a net positive mean bilateral connectivity across the cortex, which is reflective of high-quality data with a low signal-to-noise ratio, and one mouse was determined to have a mean negative bilateral measure and was discarded.

Data cleaning was also performed by discarding all frames with a standard deviation greater than 1 in the global variance of the temporal derivatives in the oxyhemoglobin signal within the 0.4-4.0 Hz band, a method adapted from Sherafati et al^50^. All calcium functional connectivity analysis was performed in the 0.4-4.0 Hz delta band. Between-group comparisons were performed as unpaired t-tests unless noted otherwise. Seed-seed analyses were performed according to the methods of White et al.^27^, in which the fluctuations of signals across time from clusters of pixels within functional regions were compared with each to determine their Pearson correlation coefficient. Homotopic contralateral (bilateral) connectivity analyses were also performed by calculating the correlation of each pixel with its counterpart reflected across the brain midline, in the contralateral hemisphere. Non-brain pixels were masked out for each mouse before analysis, and the group mask used in all bilateral and node degree analyses reflects all pixels shared by at least 149 of the total 246 imaging sessions, with those not containing data from any one pixel excluded from that pixel’s analysis. A false discovery rate multiple comparison correction was performed on all seed-seed matrix analyses, and a cluster-based correction^49^ was performed on all homotopic contralateral maps to determine significance. Time-by-genotype interactions were derived using repeated measures (rm)ANOVA analysis with Greenhouse-Geier correction for violations of sphericity, for imaging data at both P35 and their sex-specific adult time point. To determine the overall connectivity of a given pixel, we performed node degree analyses by calculating the number of other pixels with which a pixel shares a functional connection with Pearson z(r) greater than 0.3, thereby calculating the number of strong positive correlations each pixel shared with the other pixels of the brain. Node degree was calculated for every pixel within the brain field-of-view. Scatter plots of pixel-wise age-by-time data (Figures 2E, 3E, 5E, and 6E) were produced by taking the mean value of each mouse’s bilateral or node degree measures across all pixels found to have a significant time-by-genotype interaction after multiple comparison correction, which is illustrated in panel D of each respective figure. Female bilateral age-by-genotype graphs were produced by averaging all cortical pixels because no pixels survived a multiple comparison correction of the female data.

Paxinos region keys and outlines on figures were based upon the mouse brain atlas produced by Franklin and Paxinos^51^. For rotarod performance data, a main effect of genotype was calculated by using fitrm and anova functions in MATLAB to establish the significant effect of genotype, then post hoc comparisons were performed amongst the three groups (Mecp2 affected, rescue, and WT) using the anova1 and multcompare functions at each time point, for each sex. Critical alpha level was .05 unless otherwise noted.

## Author Contributions

Conceptualization, R.M.R., J.P.C. and J.D.D.; Methodology, R.M.R., J.P.C., J.D.D., A.Y. and S.E.M.; Software, L.M.B.; Formal Analysis, R.M.R. and A.Y.; Investigation, R.M.R., A.Y., S.C., S.H.G., A.R.B., R.G.S., L.L.; Writing - Original Draft, R.M.R., J.P.C., J.D.D. and A.Y.; Writing - Review & Editing, R.M.R., A.Y., S.C., S.H.G., A.R.B., L.M.B., R.G.S., L.L., S.E.M., J.P.C.., and J.D.D.; Funding Acquisition, R.M.R., J.P.C. and J.D.D.; Supervision, J.P.C. and J.D.D.

## Supplementary Information

**Figure S1:**
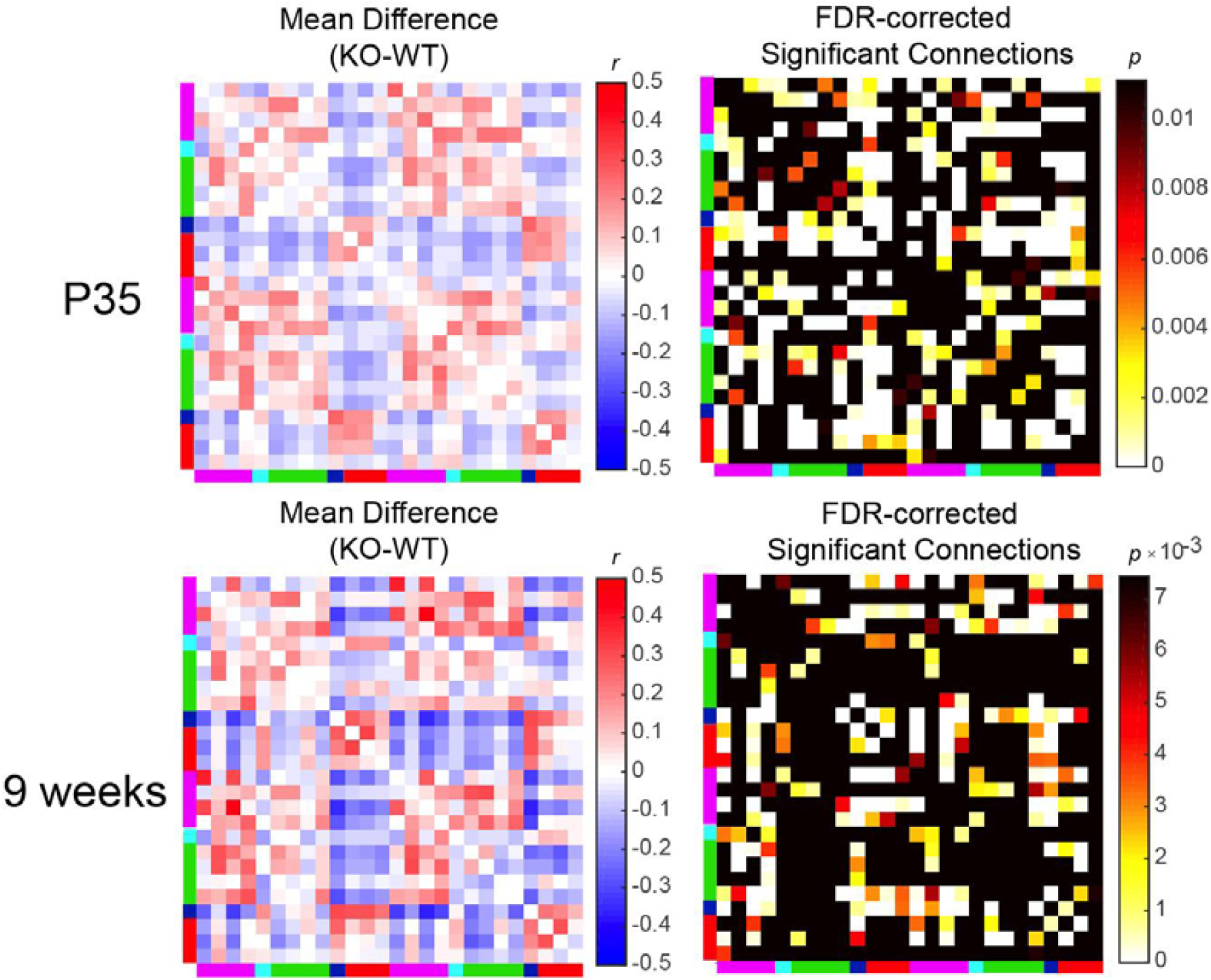
**Mean difference in connection strength and *p*-statistic matrices of Mecp2 KO males and WT controls.**

**Figure S2:**
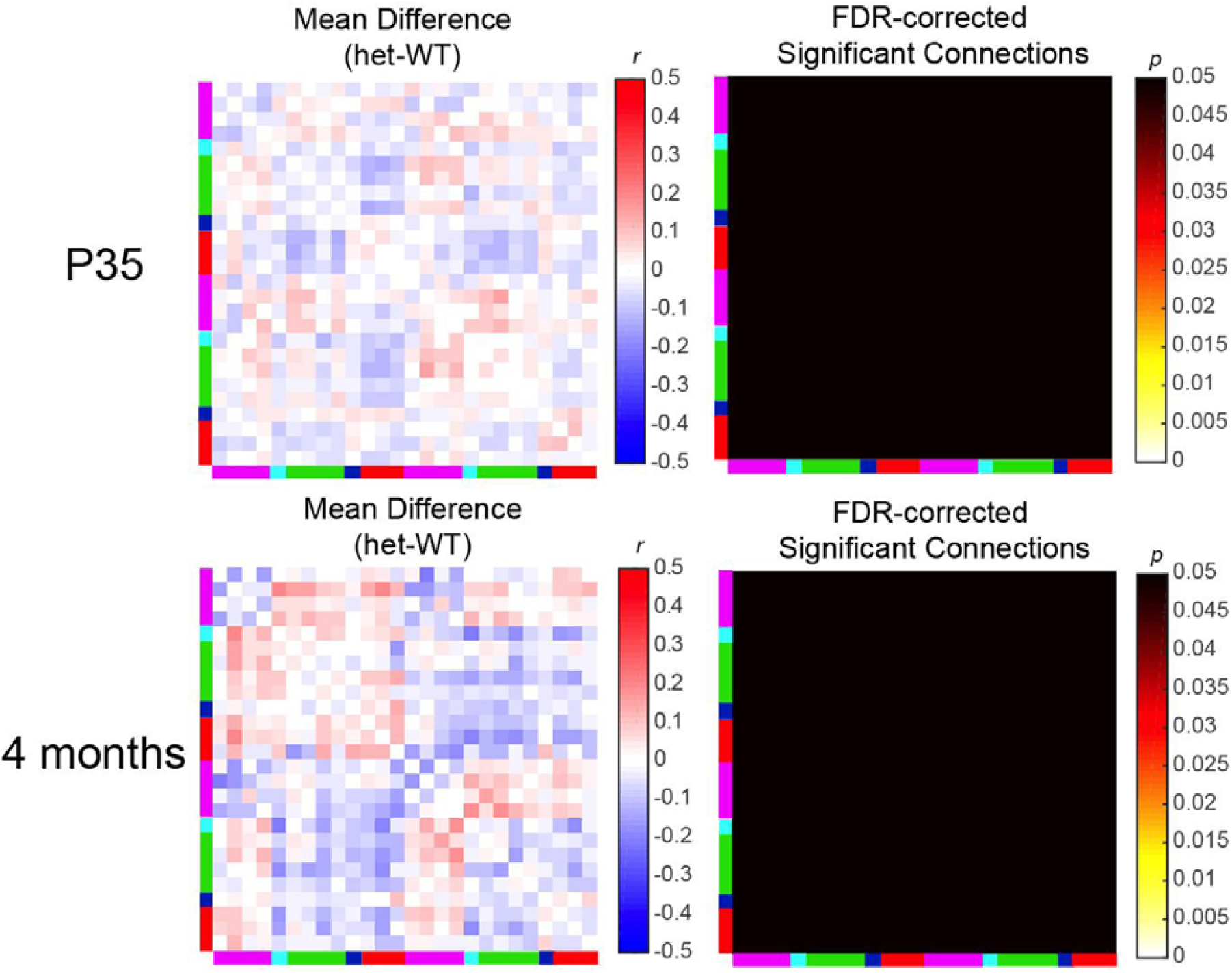
**Mean difference in connection strength and p-statistic matrices of Mecp2 heterozygous females and WT controls.**

**Figure S3:**
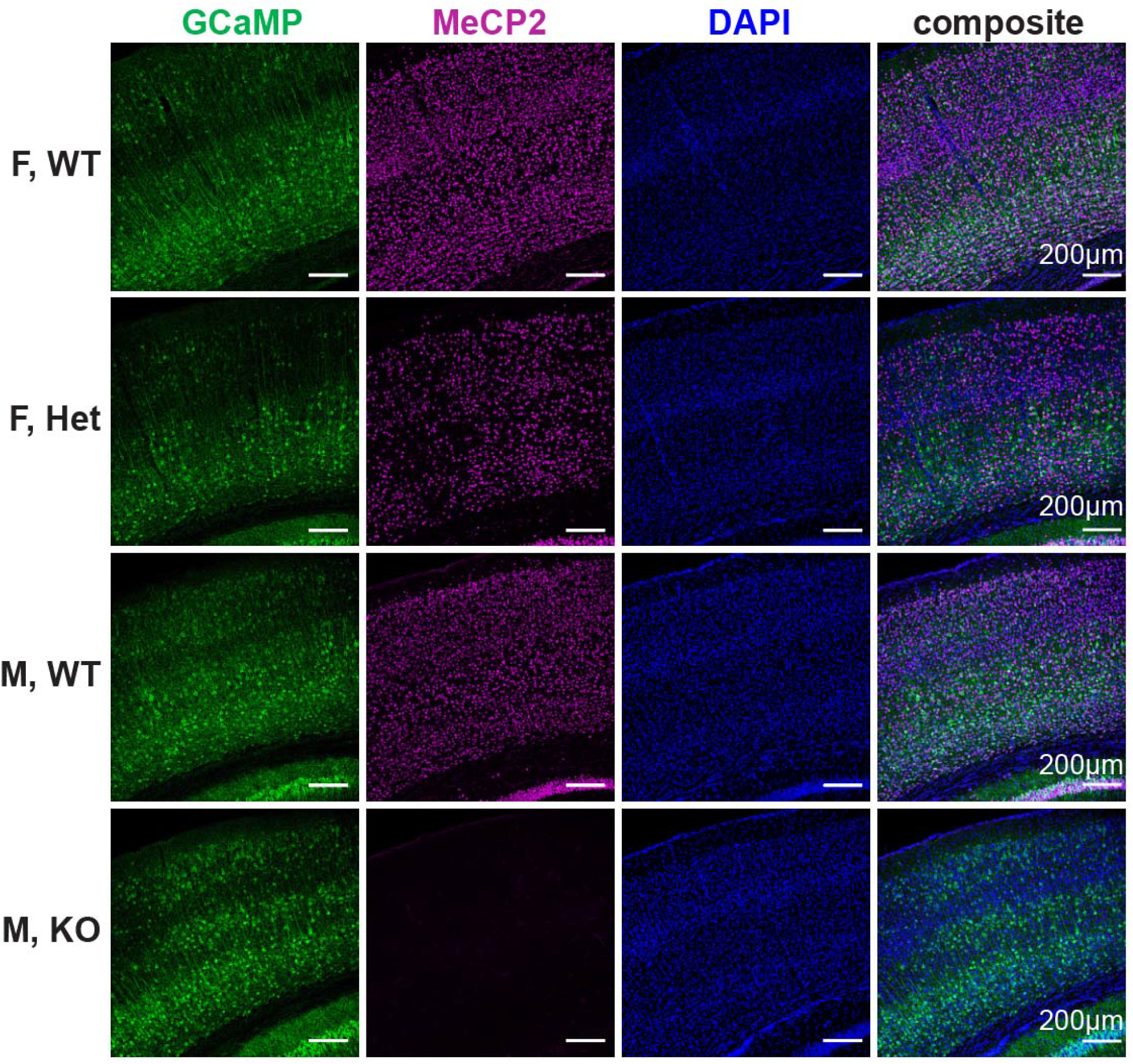
Immunofluorescence images of Mecp2 mice and WT controls confirm expected MeCP2 protein levels. Representative somatosensory cortical immunofluorescence images from male and female Mecp2 mice and WT controls, with GCaMP (calcium indicator), MeCP2, and DAPI (nuclei) stains. Het females display approximately 50% *Mecp2* expression compared to WT’s, as expected due to mosaicism linked to X-inactivation. Male KO mice display a lack of *Mecp2* expression as expected.

**Figure S4:**
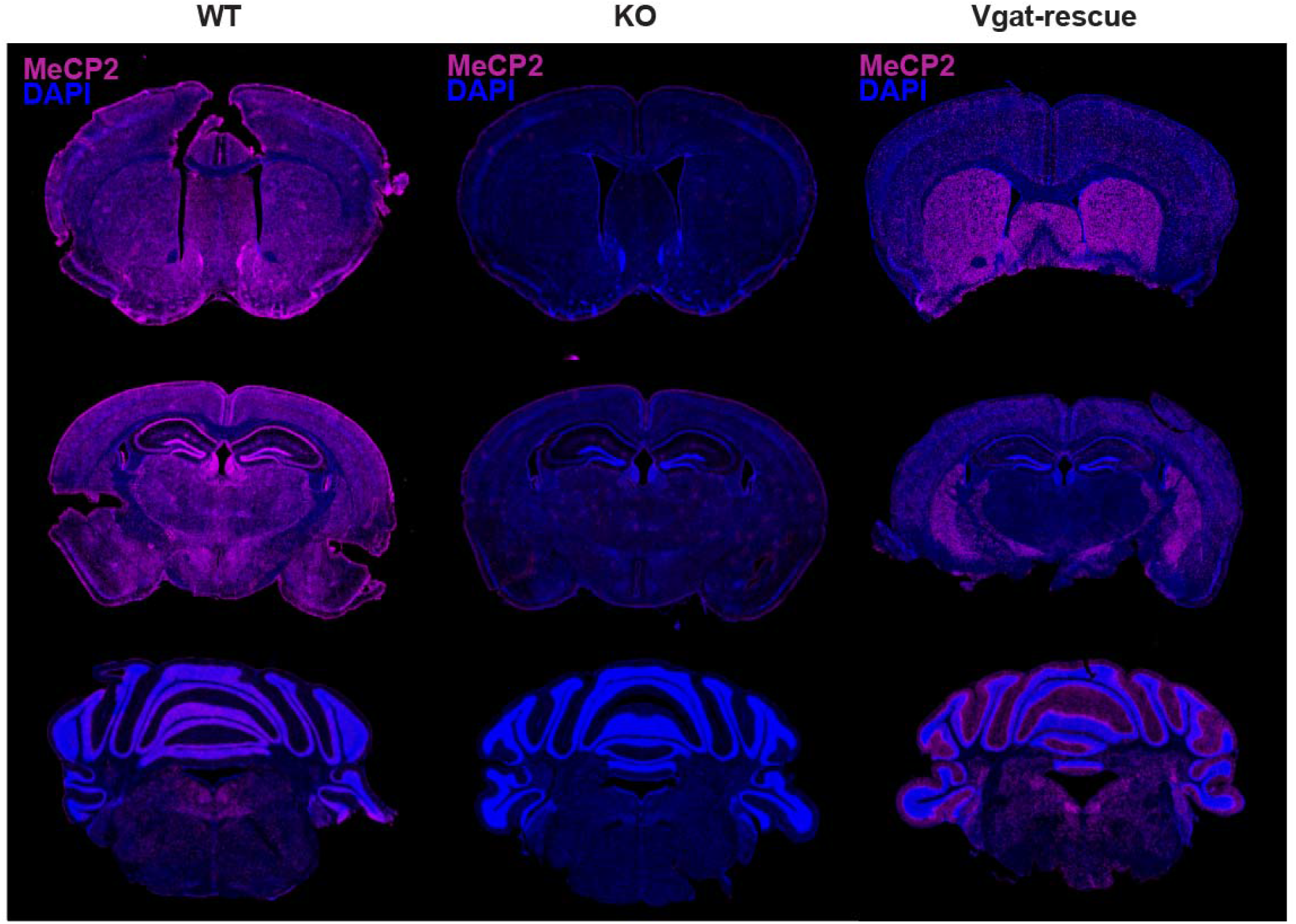
Immunofluorescence images of Mecp2 mice, WT controls, and GABAergic-specific Mecp2 rescues confirm expected MeCP2 protein levels. Representative cortical immunofluorescence images from male and female Mecp2 mice and WT controls, with MeCP2 and DAPI (nuclei) stains. KO males display a lack of *Mecp2* expression in all areas of the brain while Vgat-rescue males show stronger expression in various cortical and subcortical regions.

**Figure S5.**
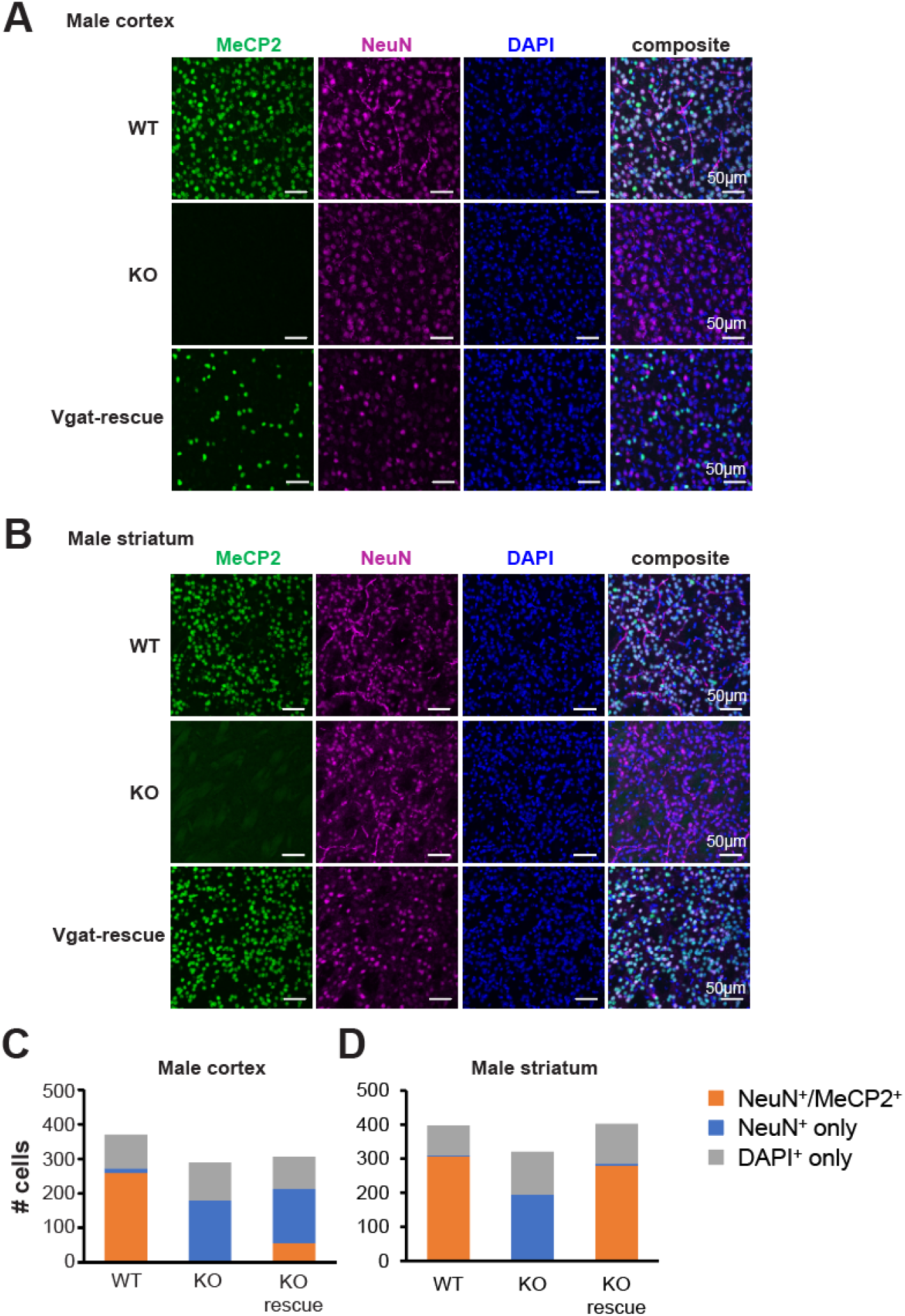
*Mecp2* expression in male KO, rescue and WT mice displays expected expression levels in cortex and striatum. Representative images show results of immunostaining for MeCP2 (green), NeuN (neuronal, purple), and DAPI (nuclei, blue) in the cortex (A) and striatum (B). Analysis of MeCP2 and NeuN colocalization shows a subtle rescue in the cortex, and a strong rescue in the striatum. (C) Summary data show the number of counted double-positive MeCP2 and NeuN neurons in the cortex, where approximately 10% of neurons are GABAergic. (D) Summary data shows nearly complete rescue in the striatum, where 90% of neurons are GABAergic. In both regions, no MeCP2 was detected in KO animals.

**Figure S6.**
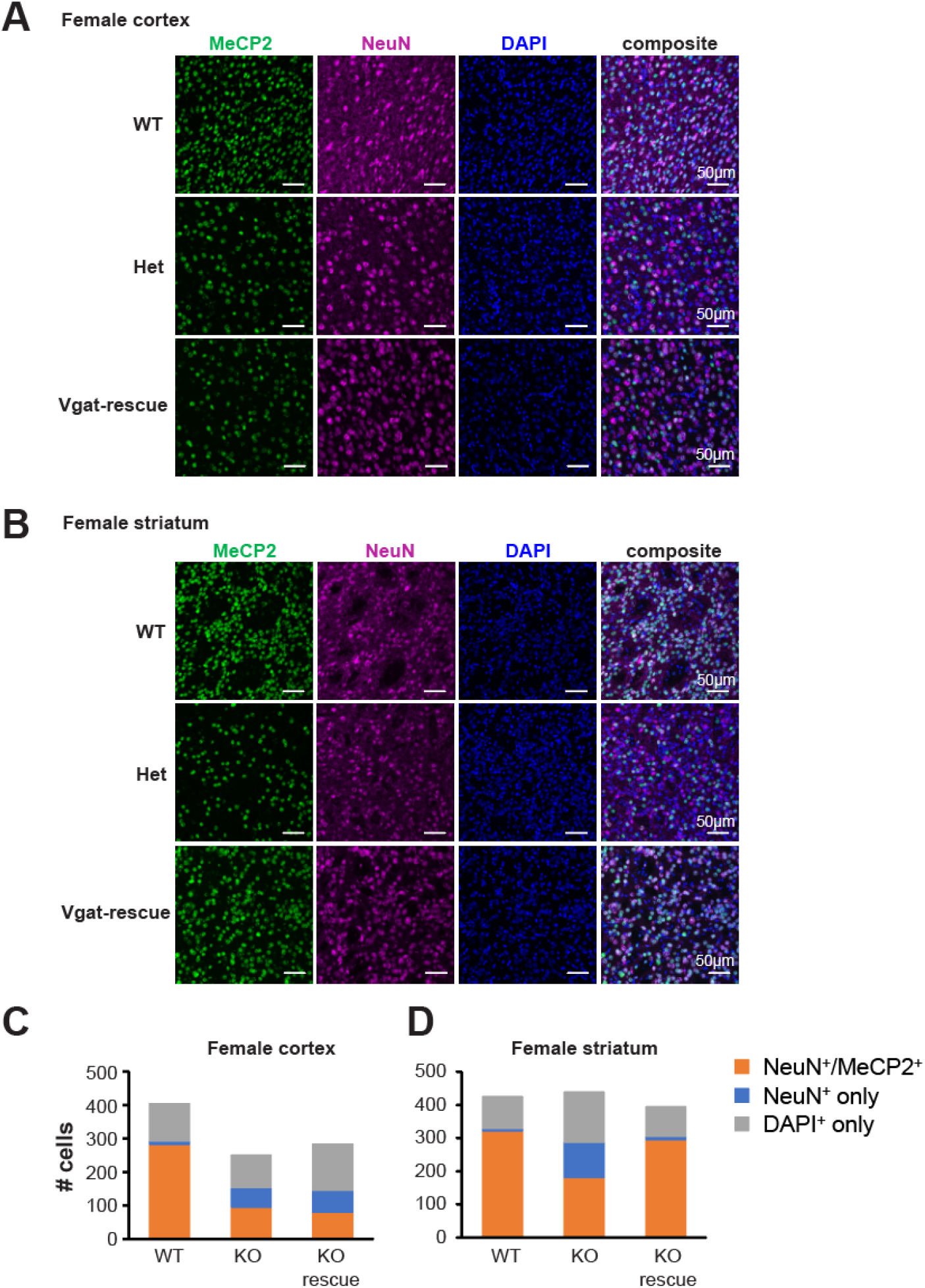
*Mecp2* expression in female heterozygous, rescue and WT mice displays expected expression levels in cortex and striatum. Representative images show results of immunostaining for MeCP2 (green), NeuN (neuronal, purple), and DAPI (nuclei, blue) in the cortex (A) and striatum (B). Females were expected to display a mosaic rescue where approximately 50% of GABAergic cells still express *Mecp2* in affected heterozygous mice due to X-inactivation, and a rescue would recover expression in the remaining 50% of the GABAergic population. Summary data show the number of counted double-positive MeCP2 and NeuN neurons cells in the cortex (C) and striatum (D). Analysis of MeCP2 and NeuN colocalization demonstrates the expected degree of MeCP2 rescue in the cortex and striatum.

**Figure S7.**
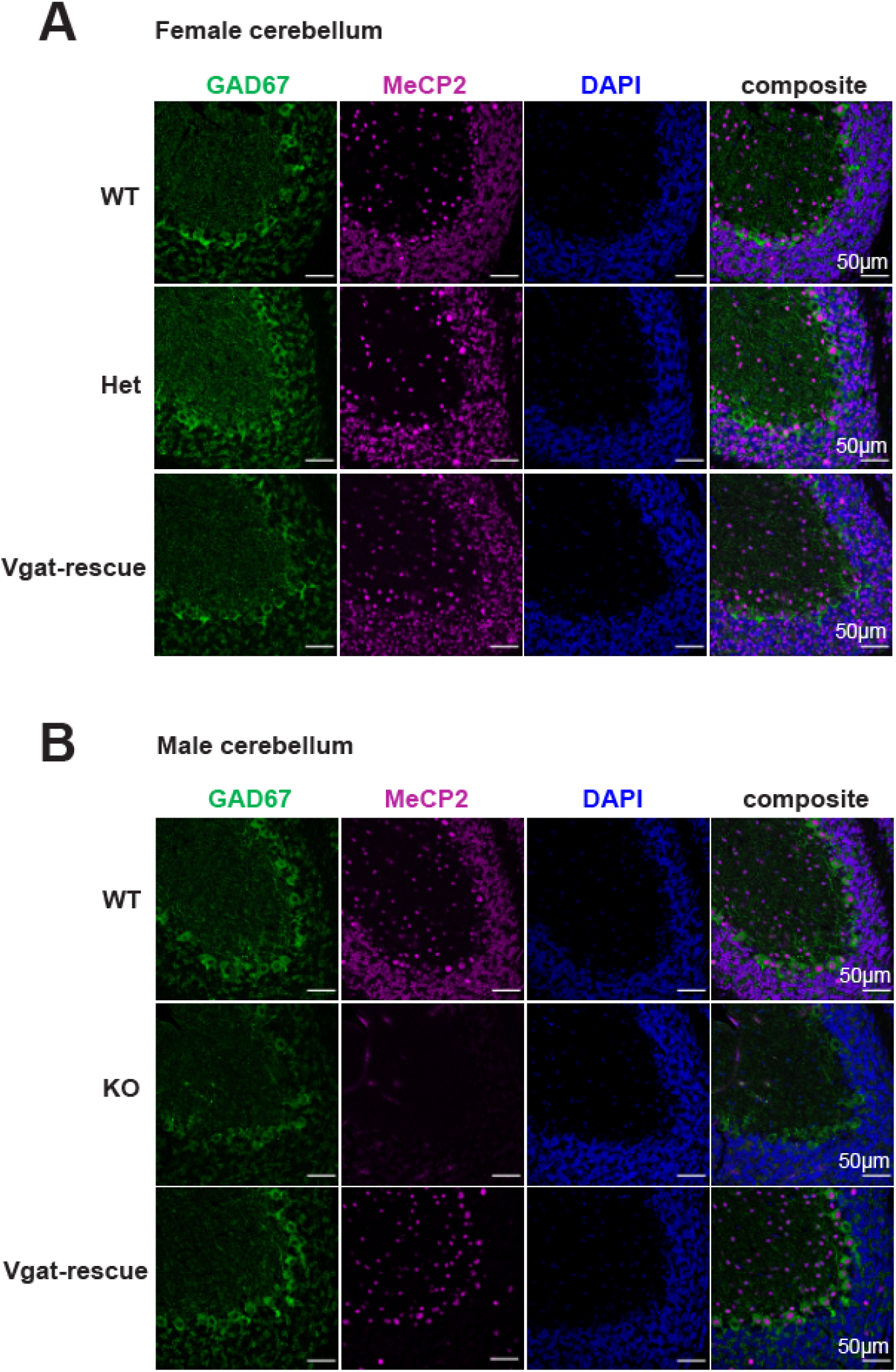
*Mecp2* expression in Mecp2 affected, rescue, and WT control mice displays expected expression levels in cerebellum. Representative images showing results of immunostaining for GAD67 (GABAergic, green), MeCP2 (purple), and DAPI (nuclei, blue) in the cerebellum of (A) female and (B) male animals. Expression of MeCP2 in GAD67+ GABAergic neurons in the Purkinje layer is reduced in hets and partially rescued in the Vgat-rescue. No *Mecp2* expression was detected in male KO animals.

